# The morphology and evolution of chondrichthyan cranial muscles: a digital dissection of the elephantfish *Callorhinchus milii* and the catshark *Scyliorhinus canicula*

**DOI:** 10.1101/2020.07.30.227132

**Authors:** Richard P. Dearden, Rohan Mansuit, Antoine Cuckovic, Anthony Herrel, Dominique Didier, Paul Tafforeau, Alan Pradel

## Abstract

The anatomy of sharks, rays, and chimaeras (chondrichthyans) is crucial to understanding the evolution of the cranial system in vertebrates, due to their position as the sister group to bony fishes (osteichthyans). Strikingly different arrangements of the head in the two constituent chondrichthyan groups – holocephalans and elasmobranchs – have played a pivotal role in the formation of evolutionary hypotheses targeting major cranial structures such as the jaws and pharynx. However, despite the advent of digital dissections as a means of easily visualizing and sharing the results of anatomical studies in three dimensions, information on the musculoskeletal systems of the chondrichthyan head remains largely limited to traditional accounts, many of which are at least a century old. Here we use synchrotron tomography acquire 3D data which we used to carry out a digital dissection of a holocephalan and an elasmobranch widely used as model species: the elephantfish, *Callorhinchus milii*, and the small-spotted catshark*, Scyliorhinus canicula*. We describe and figure the skeletal anatomy of the head, labial, mandibular, hyoid, and branchial cartilages in both taxa as well as the muscles of the head and pharynx. We make new observations, particularly regarding the branchial musculature of *Callorhinchus*, revealing several previously unreported or previously ambiguous structures. Finally, we review what is known about the evolution of chondrichthyan cranial muscles from their fossil record and discuss the implications for muscle homology and evolution, broadly concluding that the holocephalan pharynx is likely derived from a more elasmobranch-like form. This dataset has great potential as a resource, particularly for researchers using these model species for zoological research, functional morphologists requiring models of musculature and skeletons, as well as for palaeontologists seeking comparative models for extinct taxa.

## Introduction

Cartilaginous fishes (Chondrichthyes) comprise only a small fraction of total jawed vertebrate diversity (Nelson et al., 2016) but are key to understanding the evolution of jawed vertebrates. As the sister-group to the more diverse and disparate bony fishes (ray-finned fishes, lobe-finned fishes, and tetrapods) their anatomy and physiology provides a valuable outgroup comparison which, probably incorrectly, has often been held to represent primitive states” for jawed vertebrates as a “whole. The greatly divergent cranial morphologies displayed by the two constituent sister groups of Chondrichthyes – elasmobranchs and holocephalans - have themselves led to much debate, both over chondrichthyan origins and those of jawed vertebrates more broadly. For these reasons, anatomists, embryologists, and physiologists have intensively studied chondrichthyan anatomy over the last two centuries. Recently, tomographic methods have allowed the advent of digital “dissections”, where an organism s anatomy can be non-destructively visualized and communicated ‘in the form of interactive datasets. These studies have run the gamut of mammals (Cox and Faulkes, 2014; Sharp and Trusler, 2015), archosaurs (Klinkhamer et al., 2017; Lautenschlager et al., 2014), lissamphibians (Porro and Richards, 2017), and actinopterygians (Brocklehurst et al., 2019). Although aspects of cartilaginous fish anatomy have been examined (Camp et al., 2017; Denton et al., 2018; Tomita et al., 2018), three-dimensional information on the cranial musculoskeletal system is limited.

We aim to address this with a digital dissection of the hard tissues and musculature of two representatives of the Chondrichthyes: *Callorhinchus milii* Bory de Saint-Vincent 1823, a holocephalan, and *Scyliorhinus canicula* Linnaeus 1758, an elasmobranch. *Callorhinchus milii* is a callorhinchid, the sister-group to all other holocephalans (Inoue et al., 2010; Licht et al., 2012) and historically one of the best known holocephalans due to its being one of only two species to inhabit shallow, nearshore waters (Didier, 1995). As a result, the musculature of the genus has been described several times (Edgeworth, 1935; Kesteven, 1933; Luther, 1909a; Shann, 1919), most recently with Didier (1995) providing a detailed overview of the anatomy and systematics of holocephalans, including *Callorhinchus*. *Scyliorhinus canicula* is a scyliorhinid, a carcharhiniform galeomorph elasmobranch. Because of the accessibility of adults, eggs, and embryos to European researchers (it is abundant in nearshore habitats in the northeastern Atlantic) the species has featured heavily in embryological and physiological studies of chondrichthyans (Coolen et al., 2008; de Beer, 1931; Hughes and Ballintijn, 1965; Oulion et al., 2011; Reif, 1980). While accounts exist of its gross anatomy (Allis, 1917; Edgeworth, 1935; Luther, 1909b; Nakaya, 1975; Ridewood, 1899; Soares and Carvalho, 2013), detailed and fully illustrated accounts are surprisingly rare in comparison to the similarly common *Squalus*.

Here we use a synchrotron tomographic dataset to provide accounts of anatomy of the cartilages and musculature of the head in *Callorhinchus milii* and *Scyliorhinus canicula*. We examine dissection-based reports of muscle anatomy in the light of our reconstructed models. By combining this information with what is known about fossil taxa, we assess scenarios of morphological evolution of chondrichthyan cranial muscles. More broadly, we aim to provide the research community with valuable three-dimensional data on the anatomy of these taxa.

## Methods

Both specimens described here are the same as those used in Pradel et al. (2013) and full specimen acquisition and scanning details can be found therein. In brief, both specimens were scanned on beamline ID19 of the European Synchrotron Radiation Facility (ESRF). The specimen of *Callorhinchus milii* is a female at embryonic stage 36 (Didier et al., 1998), the stage immediately prior to hatching, and was originally collected from the Marlborough Sounds, New Zealand. It was scanned in a 75% ethanol solution at a voxel size of 30 microns, and a single distance phase retrieval process was used to gain differential contrast of the specimen’s tissues. The *Scyliorhinus canicula* specimen is a juvenile, reared at the Laboratoire Evolution, Génome et Spéciation, UPR 9034 CNRS, Gif-Sur-Yvette, France, which had reached the stage of independent feeding when humanely killed. It was scanned in a 100% ethanol solution using a holotomographic approach at a voxel size of 7.45 microns.

Volumes for both specimens were reconstructed using the ESRF software PyHST. Segmentations of the 3D data were carried out using Mimics versions 15-21 (Materialise). Images of the resulting three-dimensional models were created using Blender v2.80 (blender.org). Unfortunately, the resolution of the scans was not sufficient to observe innervation in most cases, so this is done throughout the text with reference to the literature.

## Results

The cranial skeletons of *Callorhinchus milii* and *Scyliorhinus canicula* have been described before in detail, but below we provide a brief description of the neurocranial and pharyngeal skeleton to supplement our account of the muscles. We also describe the muscles’ attachments and innervation. Where necessary we note disagreements about precise accounts of the innervation of the cranial muscles. The supporting information includes 3D files of all structures described (Dearden *et al*. 2020, SI1-3).

### Callorhinchus milii

#### Cranial Cartilages

The head skeleton of *Callorhinchus* comprises the neurocranium (Fig. 1), to which the palatoquadrates are fused, the mandible, formed from the two medially fused Meckel’s cartilages, a series of six paired labial cartilages surrounding the mouth, a non-suspensory hyoid arch, and five branchial arches (Fig. 2). Like all other living holocephalans the head skeleton is antero-posteriorly compact: the lower jaw extends posteriorly only as far as the back of the orbits, and all other pharyngeal cartilages are located ventral to the neurocranium. The whole arrangement is posteriorly bounded by ventrally joined scapulocoracoids.

**Figure 1.**
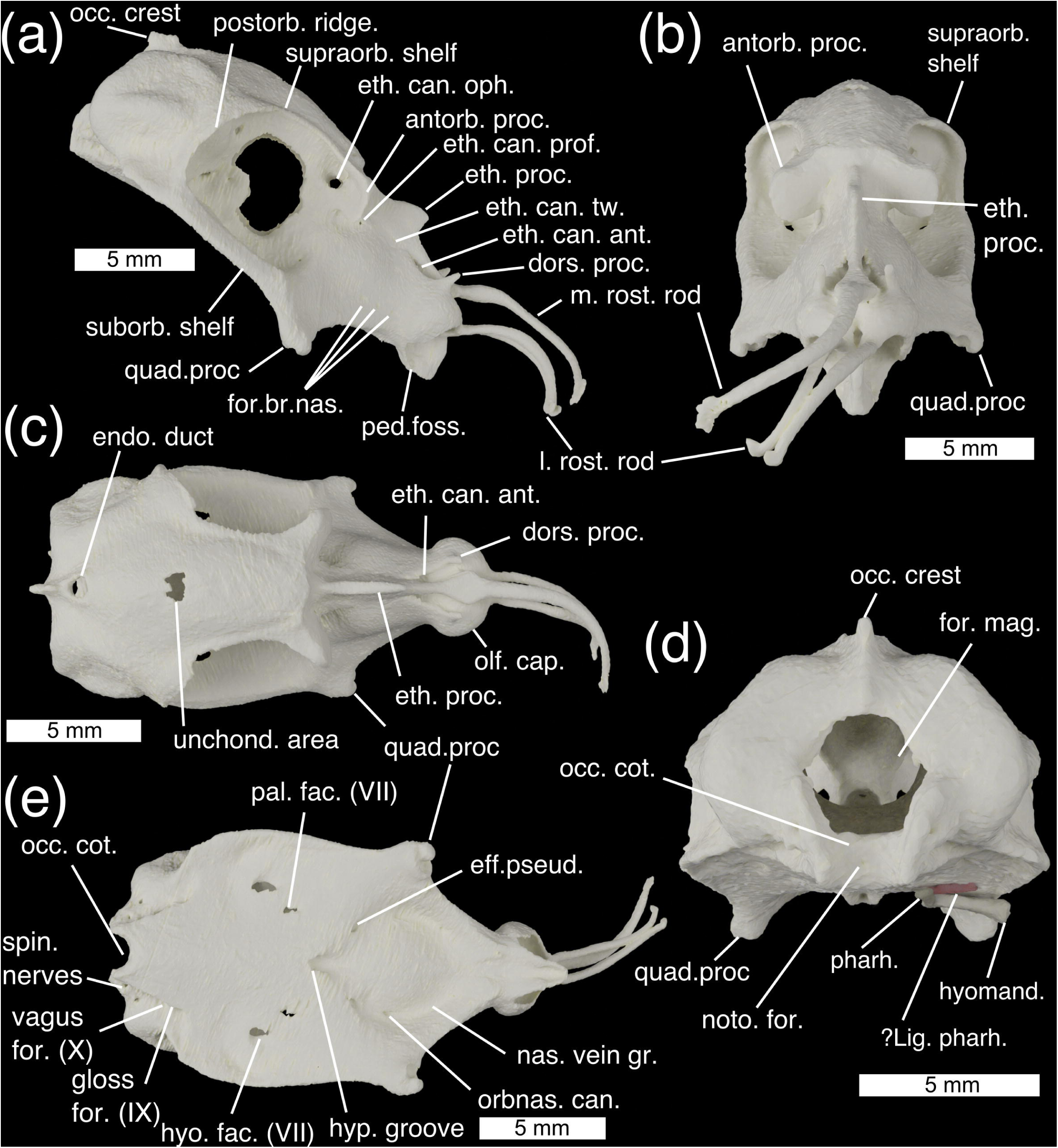
The neurocranium of *Callorhinchus milii* in (a) lateral, (b) anterior, (c) dorsal, (d) posterior, and (e) ventral views. Abbreviations: antorb. proc., antorbital process; dors. proc. dorsal process; eff. pseud., foramen for the efferent pseudobranchial artery; endo. duct, foramen for endolymphatic duct; eth. can. Ant, anterior ethmoid canal foramina eth. can. oph, entry into ethmoid canal for superficial ophthalmic complex; eth. can. prof, entry into ethmoid canal for profundus nerve; eth. can. tw., exit from ethmoid canal for twigs of the superficial ophthalmic complex/profundus; eth. proc., ethmoid process; for. mag.; foramen magnum; for. br. nas., foraminae for branches of the nasal vein; gloss for. (IX), foramen for glossopharyngeal (IX) nerve; hyo. fac. (VII), foramen for hyomandibular branch of the facial (VII) nerve; hyomand., hyomandibula; hyp. groove., hypophyseal groove; l. rost. rod, lateral rostral rod; ?*lig. pharh.*, *?ligamentum pharyngohyoideus*; m. rost. rod, median rostral rod; nas. vein gr., groove for the nasal vein; noto. for., notochord foramen; occ. crest, occipital crest; occ. cot., occipital cotylus; olf. cap., olfactory capsules; orbnas. can., foramen for the orbitonasal canal; pal. fac. (VII), foramen for palatine branch of the facial (VII) nerve; ped. foss., fossa for pedicular cartilage; pharh, pharyngohyal; postorb. ridge, postorbital ridge; quad. proc., quadrate process; spin. nerves, foraminae for anterior spinal nerves; suborb. shelf, suborbital shelf; supraorb. shelf, supraorbital shelf; unchond. area, incompletely chondrified area; vagus for. (X), foramen for vagus (X) nerve

**Figure 2.**
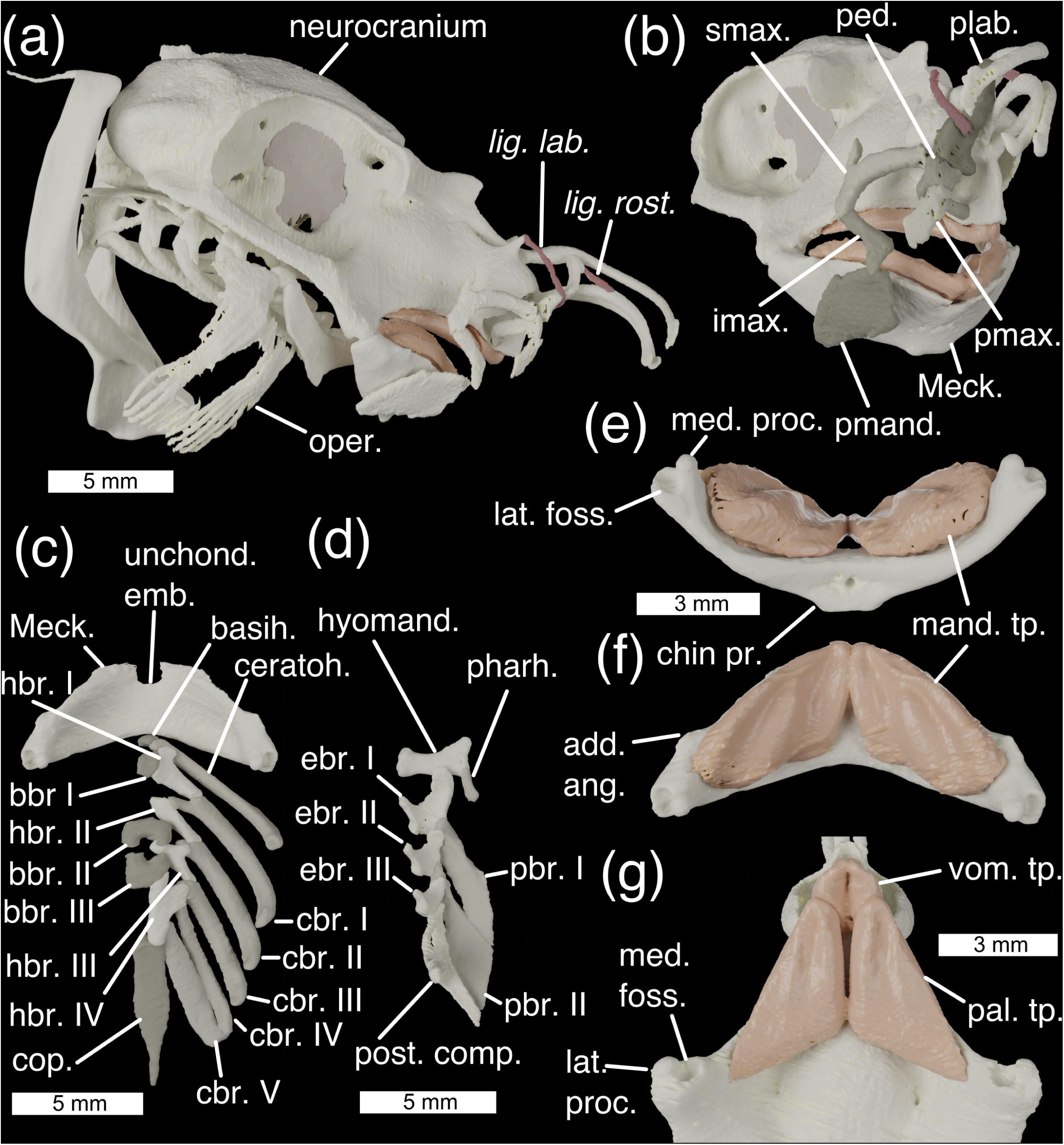
The cranial skeleton of *Callorhinchus milii* (a) complete skeleton in lateral view, (b) antero-lateral view showing labial cartilages, (c) ventral pharyngeal skeleton in dorsal view, (d) dorsal pharyngeal skeleton in ventral view, lower jaw in ventral (e) and dorsal (f) view, and (g) palate in ventral view. Colours: cream, cartilage; beige, sphenoptic membrane; red, ligaments. Abbreviations: add. ang., adductor mandibulae angle; bbr., basibranchials; basih., basihyal; cbr., ceratobranchial; ceratoh, ceratohyal; chin pr., chin process; cop., basibranchial copula; ebr, epibranchial; hyomand., hyomandibula; hbr., hypobranchial; imax., inferior maxillary cartilage; lat. foss., lateral fossa of Meckel’s cartilage; lat. proc., lateral process of quadrate; *lig. lab., ligamentum labialis*; *lig*. *rost*., *ligamentum rostralis*; mand. tp., mandibular toothplates; Meck., Meckel’s cartilage; oper., opercular cartilage; med. foss, medial fossa of quadrate; med. proc., medial process of Meckel’s cartilage; pal. tp. palatine toothplates; ped. pedicular cartilage; pharh, pharyngohyal; plab., prelabial cartilage; pmand. premandibular cartilage; pbr., pharyngobranchial; pmax., premaxillary cartilage; post. comp., posterior epibranchial/pharyngobranchial complex; smax., superior maxillary cartilage; unchond. emb., unchondrified embayment; vom. tp., vomerine toothplates

The **neurocranium** of *Callorhinchus milii* is tall, with an extensive rostrum, enlarged orbits, and a laterally broad otic region (Fig. 1). The olfactory capsules (Fig. 1c; olf. cap.) take the form of two rounded, ventrally open bulbs, closely set at the extreme anterior end of the neurocranium. A short dorsal process (Fig. 1a, c; dors. proc.) projects from the apex of each capsule. Between the olfactory capsules three long rostral rods project anteriorly to support the animal’s “trunk”: one median rod along the midline (Fig. 1a, b; m. rost. rod) and a pair of lateral rods (Fig. 1a, b; l. rost. rod). Below them a beak projects ventrally, forming the anterior end of the mouth’s roof, and carrying the vomerine toothplates (Fig. 2g; vom. tp.). At the very base of the beak’s lateral sides are a pair of small fossae with which the pedicular cartilages articulate (Fig. 1a, ped. foss.). The ethmoid region is long, with steeply sloping sides separated at the midline by the ethmoid crest which has a marked ethmoid process at its middle (Fig. 1a, b, c, eth. proc.). A pair of foraminae in the orbits form the entrance to the ethmoid canal for the superficial ophthalmic complex (Fig. 1a; eth. can. oph.), which runs anteriorly through the midline length of the ethmoid region. About two fifths of the way along the canal’s length it is punctured by a lateral foramen for the entry of the profundus (V_1_) nerve (Fig. 1a; eth. can. prof.), anterior to this several small foraminae along its length allow twigs of the superficial ophthalmic complex + profundus to exit onto the ethmoid surface (Fig. 1a; eth. can. tw.). The canal opens anteriorly through a pair of teardrop-shaped foraminae posterior to the nasal capsules (Fig. 1a, c; eth. can. ant.). On the ventral slope of the ethmoid region, a row of three foraminae provide passage for branches of the nasal vein behind the nasal capsules (Fig. 1a; for. br. nas.). At the postero-ventral corner of the ethmoid region are a pair of stout quadrate processes (Fig. 1a, b, e; quad. proc.), which flare laterally to meet the Meckelian cartilages (Fig. 2b; Meck.).

The orbits in *Callorhinchus* are very large, occupying the neurocranium’s full height and about two fifths of its length. Anteriorly they are bounded by laterally projecting antorbital processes (Fig. 1a, b; antorb. proc.) and a preorbital fascia (Didier, 1995). The inner orbital wall is formed by an extensive sphenoptic membrane (Figs. 2, 3). Posterior to the antorbital processes the roof of the neurocranium pinches in laterally, meaning that there only a narrow supraorbital shelf (Fig. 1a, b; supraorb. shelf) to the orbits, before expanding posteriorly into the postorbital ridge (Fig. 1a; postorb. ridge), which curves ventrally to form the rear wall of the orbit. Ventrally the orbits are bounded by a broad, flat suborbital shelf (Fig. 1a; suborb. shelf), which at its lateral extent broadens into a marked ridge. Just above the level of the suborbital shelf is a large foramen in the sphenoptic membrane for the optic (II) nerve (Fig. 3a, b; opt. (II) op.), and immediately posteriorly to this is a small opening for the central retinal (optic) artery (Fig. 3a, b; ret. art.). Antero-ventrally is a small orbitonasal canal foramen for the nasal vein (Fig. 3a; orbnas. can.). The superficial ophthalmic complex (V + anterodorsal lateral line) enters the orbit through a dorso-posterior foramen (Fig. 3a, b; sup. oph. for.), and then exits, entering the ethmoid canal, via a large ophthalmic foramen in the antero-dorsal part of the orbit (Fig. 3a; oph. for.). In the posteroventral corner of the orbit is a large foramen through which the trigeminal (V) and facial (VII) nerves enter the orbit (Fig. 3a, b; for. V + VII). Two foraminae in the ventral part of the orbit provide exits for the hyomandibular (Figs. 1e, 3b; hyo. fac. (VII)) and palatine (Figs. 1e, 3b; pal. fac. (VII)) branches of the facial nerve onto the neurocranial floor. The orbital artery also enters the orbit through the palatine foramen.

**Figure 3.**
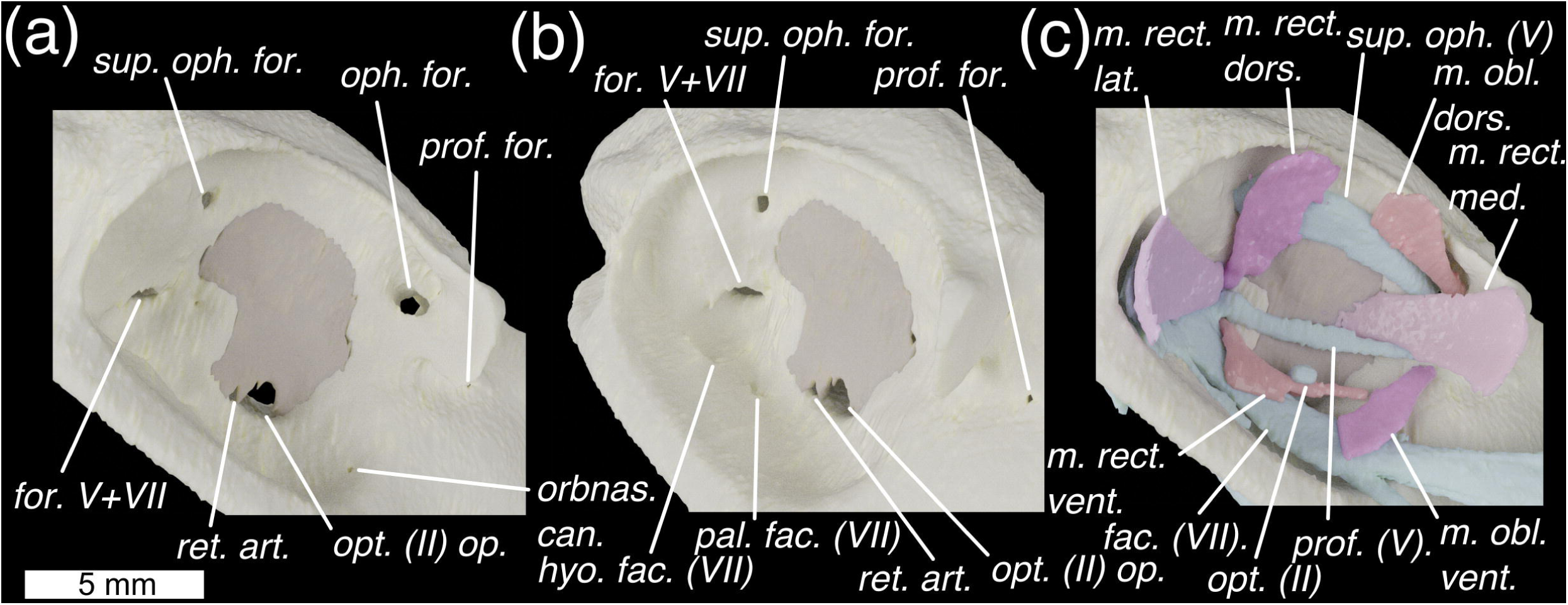
The orbit of *Callorhinchus milii* shown in lateral view with skeleton in (a) lateral and (b) antero-lateral view and (c)lateral view with external optic muscles and cranial nerves. Colours as in Fig. 3 with light-blue for cranial nerves. Abbreviations: fac. (VII), facial (VII) nerve; hyo. fac. (VII), foramen for hyomandibular branch of the facial (VII) nerve; *m. obl. dors.*, *m. obliquus dorsalis*; *m. obl. vent.*, *m. obliquus ventralis*; *m. rect. dors.*, *m. rectus dorsalis*; *m. rect. lat.*, *m. rectus lateralis*; *m. rect. med.*, *m. rectus medialis*; *m. rect. vent.*, *m. rectus ventralis*; oph. for. ophthalmic foramen; opt. (II), optic nerve; opt. (II) op., optic nerve opening; orbnas. can., foramen for the orbitonasal canal; pal. fac. (VII), foramen for palatine branch of the facial (VII) nerve; prof. for., foramen for profundus; prof. (V), profundus (V) nerve; ret. art., retinal artery opening; sup. oph. (V), superficial ophthalmic complex (V+anterodorsal lateral line nerves); sup. oph. for., foramen for superficial ophthalmic complex’s entry into orbit; for. V+VII, foramen for entry of trigeminal (V) and facial (VII) nerves into the orbit.

Between the orbits the neurocranial roof forms a shallowly convex surface, which becomes more pronounced posteriorly. This shallow roof curves slightly ventrally, and at its apex is a small, unchondrified area (Fig. 1c; unchond. area). The roof pinches in laterally to meet the endolymphatic duct opening (Fig. 1c; endo. duct), which is large and subcircular. Posteriorly to this is an occipital crest (Fig. 1d; occ. crest), which is pronounced dorsally, before becoming lower and being interrupted by the foramen magnum. The otic capsules form two pronounced bulges on either side of the neurocranium, with the anterior, posterior, and lateral canals forming a rough triangle of ridges, dorso-anteriorly, posteriorly, and laterally. Ventral to the lateral ridge the sides of the neurocranium pinch in before expanding again to form the edge of the neurocranial floor. The foramen magnum (Fig. 1d; for. mag.) is a large circular opening about half the height of the neurocranium, ventral to which is the shallow, rectangular occipital cotylus (Fig. 1d; occ. cot.), bounded by two long, thin condyles sit on either side. A small foramen in the centre of the occipital cotylus permits entry for the notochord (Fig. 1d; noto. for.), which extends anteriorly into the *dorsum sellae*.

The ventral surface of the neurocranium is markedly broad and flat, tapering and becoming slightly convex anteriorly where it forms the roof of the mouth. The palatine toothplates (Fig. 2g; pal. tp.) sit between the vomerine toothplates and the quadrate processes. At the approximate centre of the neurocranial floor lies a deep hypophyseal groove (Fig. 1e; hyp. groove.). Postero-laterally to the hypophyseal groove are the two paired foraminae through which the hyomandibular (Figs. 1e, 3b; hyo. fac. (VII)) and palatine (Figs. 1e, 3b; pal. fac. (VII)) branches of the facial nerve (VII) exit the orbit. Antero-laterally to the hypophyseal groove are a pair of foraminae through which the efferent pseudobranchial arteries enter the braincase (Fig. 1e; eff. pseud.). Anteriorly to these are paired orbitonasal canals through which the nasal veins pass into the orbit (Fig. 1e; orbnas. can.), which mark the posterior end of paired grooves carrying the nasal veins over the neurocranial floor from the nasal capsule to the orbits (Fig. 1e; nas. vein gr.). Posteriorly, either side of the neurocranial floor tapers medially and joins the otic region, while the central floor continues posteriorly to form the base of the occiput. Just dorsally to this is a row of foraminae for the glossopharyngeal (IX) nerve (Fig. 1e; gloss for. (IX)), vagus (X) nerve (Fig. 1e; vagus for. (X)), and anterior spinal nerves (Fig. 1e; spin. nerves).

**Meckel’s cartilages** (Fig. 2b; Meck.) are fused at a medial symphysis to form a single, bow-shaped element. This element has a flat surface, the ventral face of which is shallowly convex. Shallow fossae in the dorsal surface carry the mandibular toothplates (Fig. 2e, f; mand. tp.). The ventro-posterior midline deepens into a pronounced chin process (Fig. 2e, chin pr.) onto which the *m. mandibulohyoidei* attach. At the anterior midline is an unchondrified embayment (Fig. 2b, unchond. emb.). The articular regions of the mandibular cartilages are positioned at their extreme posterior ends, and as in other chondrichthyans, are double-articulating. A lateral process (Fig. 2g; lat. proc.) and a medial fossa (Fig. 2g; med. foss.) on the quadrate process articulate with a medial process (Fig. 2e; med. proc.) and lateral fossa (Fig. 2e; lat. foss.) on Meckel’s cartilage. Anteriorly to the articulation on each side of the Meckel’s cartilage is a small angle onto which the *m. adductor mandibulae posterior* attaches (Fig. 2f; add. ang.).

Six pairs of **labial cartilages** are present in *Callorhinchus*: the premandibular, inferior maxillary, superior maxillary, pedicular, premaxillary, and prelabial. These surround the mouth, supporting the animal’s fleshy lip, and provide the insertion surfaces for a series of muscles and tendons. The premandibular cartilages (Fig. 2b; premand.) are broad and plate-like, sitting laterally to the mandibles. Above them are the short, hockey-stick shaped inferior maxillary (Fig. 2b; imax.) cartilages. The dorsal tip of these articulate with the superior maxillary cartilages (Fig. 2b; smax.) – large, curved elements with a dorsal process. The anterior tip of this meets the back of the round head of the pedicular cartilage (Fig. 2b; ped., which posteriorly curves medially to meet a small fossa in the ethmoid region. Ventrally the head of the pedicular cartilage meets the premaxillary cartilage, a short, flat element which extends into the tissue of the upper lip (Fig. 2b; pmax.). Dorsally the head of the pedicular cartilage meets the prelabial cartilage (Fig. 2b; plab.), a gentle sigmoid that extends dorsally, and which is secured to the rostral rods by labial and rostral ligaments.

The **hyoid arch** does not articulate with the neurocranium, and comprises a basihyal and paired ceratohyals, hyomandibulae (epihyals), and pharyngohyals. The basihyal (Fig. 2c, basih.) is a very small, approximately triangular element with shallow lateral fossae where it articulates tightly with the ceratohyals. The ceratohyal (Fig. 2c, ceratoh.) is a large, curved, and flattened element, with a pronounced ventral angle. The hyomandibula (Fig. 2d, hyomand.), or epihyal, is smaller and flat with pronounced corners, and articulates with the ceratohyal at its ventral corner. At its dorsal corner it articulates with the small, ovoid pharyngohyal (Fig. 2d, pharh.). The posterior corner of the hyomandibula is in contact with the opercular cartilage (Fig. 2a, oper.). This has a broad, flat antero-dorsal corner, which divides posteriorly into a series of parallel rays divided roughly into dorsal and ventral zones. Posteriorly to the hyoid arch no branchial or extrabranchial rays are present.

The **branchial skeleton** comprises five branchial arches, lying entirely ventrally to the neurocranium. The midline floor of the pharynx is formed by a series of basibranchials (Fig. 2c, bbr.), a very small anterior one, two middle ones that between them form a box-shape, and a long, posterior basibranchial copula (Fig. 2c, cop.). Articulating with these are four paired hypobranchials (Fig. 2c; hbr). These are directed antero-medially, each contacting at least two pharyngeal arches. The first hypobranchial contacts the ceratohyal and first ceratobranchial as well as the anterior basibranchial, the second hypobranchial contacts the first and second ceratobranchials, the third hypobranchial contacts the second and third ceratobranchials and extends between the second and third basibranchials, while the fourth hypobranchial is larger than the others and contacts the third, fourth, and fifth ceratobranchials in addition to the third basibranchial and the basibranchial copula. The ceratobranchials are gently curved, with short processes to contact the hypobranchials, and the fifth ceratobranchial is slightly flattened and closely associated with the fourth. The first three branchial arches have three separate epibranchials which are flattened and square, with a ventral process that articulates with the ceratobranchials (Fig. 2c; cbr). They also have a pronounced anterior process that in the first arch, underlies the hyomandibula, and in the posterior arches contacts the anterior pharyngobranchial. Two separate pharyngobranchials (Fig. 2d; pbr.) articulate on the posterior corners of the first two epibranchials, projecting posteriorly, also contacting the anterior processes of the second and third epibranchials. They are well-developed and flat, with a dorsal groove over which the second and third efferent branchial arteries pass. At the posterior end of the dorsal branchial skeleton is a complex cartilage (Fig. 2d; post. comp.), taking the place of the fourth and fifth epibranchials, as well as the third, fourth, and fifth pharyngobranchials. This structure is roughly triangular in shape, and shallowly convex medially, with pronounced postero-ventral and postero-dorsal processes. On its antero-ventral side it contacts the fourth and fifth ceratobranchials, before projecting anteriorly into a process that contacts the postero-dorsal corner of the third epibranchial. The dorsal surface of this process is obliquely grooved for the fourth efferent branchial artery, and there is a large foramen in this anterior process through which the fifth branchial artery passes.

#### Cranial Musculature

This account follows the terminology of Didier (1995) and Anderson (2008), which itself follows Vetter (1878), Edgeworth (1935), and Shann (1919).

***M. adductor mandibulae posterior*** (Fig. 4a; *m. add. mand. post.*)

**Figure 4.**
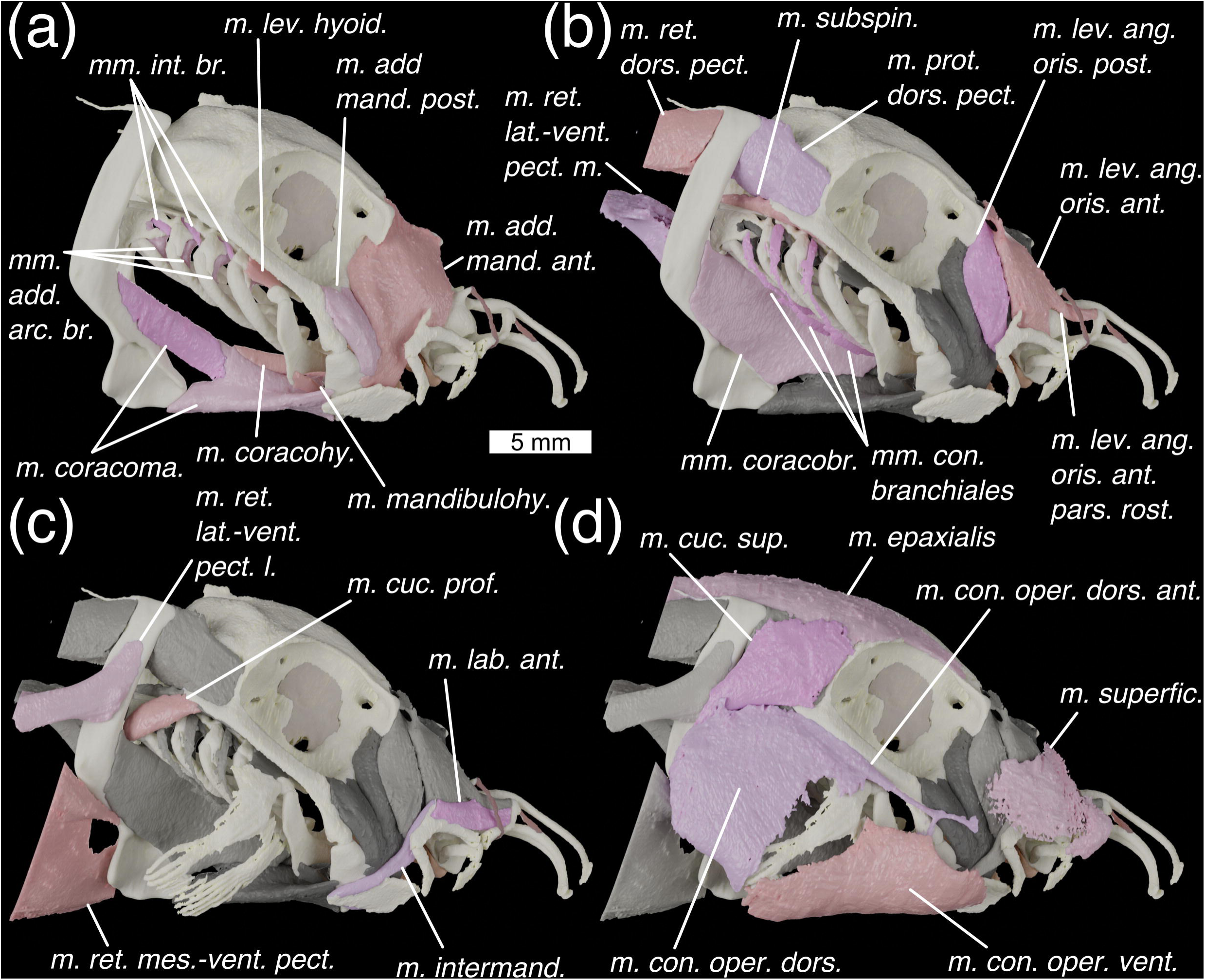
Lateral view of the head of *Callorhinchus milii* with progressively shallower muscles shown. Colours: cream, cartilage; beige, sphenoptic membrane; red, ligaments; pinks, muscles; greys, deeper muscles. Abbreviations: *mm. add. arc. br., mm. adductors arcuum branchiales; m. add. mand. ant.*, *m. adductor mandibulae anterior*; *m. add. mand. post.*, *m. adductor mandibulae posterior; mm. con. branchiales, mm. constrictors branchiales*; *m. con. oper. dors,. m. constrictor opercula dorsalis*; *m. con. oper. dors. ant., m. constrictor operculi dorsalis anterior*; *m. con. oper. vent., m. constrictor operculi ventralis; m. coracohy., mm. coracohyoideus*; *mm. coracobr., mm. coracobranchiales; m. coracoma., m. coracomandibularis* (coloured in two shades to show division)*; m. cuc. prof., m. cucullaris profundus; m. cuc. sup., m. cucullaris superficialis; mm. int. br., m. interarcuales branchiales; m. intermand.*, *m. intermandibularis*; *m. lab. ant.*, *m. labialis anterior; m. lev. ang. oris ant.*, *m. levator anguli oris anterior; m. lev. ang. oris ant. pars. rost., m. levator anguli oris anterior pars rostralis*; *m. lev. ang. oris post.*, *levator anguli oris posterior*; *m. lev. hyoid., m. levator hyoideus; m. mandibulohy., m. mandibulohyoideus; m. prot. dors. pect., m. protractor dorsalis pectoralis; m. ret. dors. pect., m. retractor dorsalis pectoralis; m. ret. lat.-vent. pect. l, m. retractor latero-ventralis pectoralis lateral; m. ret. lat.-vent. pect. m, m. retractor latero-ventralis pectoralis medial; m. ret. mes.-vent. pect., m. retractor mesio-ventralis pectoralis; m. subspin, m. subspinalis; m. superfic.*, *m. superficialis*

*Description:* A small muscle with an origin along the anterior edge of the suborbital shelf, antero-ventrally to the orbit. Didier (1995) also reports an origin on the preorbital fascia, which is not visible in our scans – if this is present it is small. The muscle lies in a shallow depression anterior to the quadrate process, anteriorly overlapping the *m. adductor mandibulae anterior,* with a surface divided into two parts by the passage of the mandibular branch of the trigeminal nerve (V_3_). It inserts over a small angle on Meckel’s cartilage (Fig. 2f; add. ang.), joining with a sheet of connective tissue slung ventrally around Meckel’s cartilage.

*Innervation:* Trigeminal (V) nerve (Didier, 1995).

*Remarks:* The *m. adductor mandibulae posterior*’s extent is variable in different holocephalan genera, and is relatively reduced in *Rhinochimaera* and *Chimaera* (Didier, 1995).

***M. adductor mandibulae anterior*** (Fig. 4a; *m. add. mand. ant.*)

*Description*: Larger than the *m. adductor mandibulae posterior* and lies entirely preorbitally. Its very broad origin extends from the preorbital fascia, across the medial section of the antorbital process, and along the ethmoid crest. It narrows ventrally to insert over the posterior part of Meckel s cartilage, joining the connective tissue which is slung around the lower jaw. An internal sheet of tissue divides the posterior quarter of the muscle from the anterior three-quarters.

*Innervation:* Trigeminal (V) nerve (Didier, 1995).

*Remarks:* In male holocephalans this muscle is functionally linked to the frontal tenaculum (Didier, 1995; Raikow and Swierczewski, 1975).

***M. levator anguli oris posterior*** (Fig. 4b; *m. lev. ang. oris post.*)

*Description:* This muscle has an origin on the antorbital crest and preorbital fascia. It passes ventrally over the *m. adductor mandibulae anterior,* inserting onto the connective tissue of the lip between the premandibular and inferior maxillary cartilages via a tendon.

*Innervation:* Trigeminal (V) nerve (Didier, 1995).

*Remarks:*The extent of the origin of this muscle is variable in different genera (Didier, 1995). Didier also reports an insertion on the supramaxillary cartilage – however, in our scans it appears to bypass the cartilage medially, separated from it by the *m. intermandibularis* (Fig. 4c; *m. intermand.*).

***M. levator anguli oris anterior*** (Fig. 4b; *m. lev. ang. oris ant.*)

*Description:* This muscle, has an origin on the connective tissue attaching to the antorbital crest. It inserts along the medial side of the anterior part of the superior maxillary cartilage. A bundle of fibres, the ***M. levator anguli oris anterior pars rostralis*** (Fig. 4b; *m. lev. ang. oris ant. pars. rost.*), extends anteriorly and inserts on the posterior side of the dorsal part of the prelabial cartilage. The fibres of this portion are mixed with those of the *m. levator anguli oris anterior* posteriorly and with those of the *m. labialis anterior* anteriorly.

*Innervation:* Trigeminal (V) nerve (Didier, 1995).

*Remarks:* In male holocephalans this muscle is functionally linked to the frontal tenaculum (Didier, 1995; Raikow and Swierczewski, 1975). Didier (1995) reported that the *M. levator anguli oris anterior pars rostralis* inserted on the rostral rod in *Callorhinchus.* We find instead that it inserts on the prelabial cartilage, consistent with the states that Didier reports in *Rhinochimaera* and *Chimaera*. Anderson (2008) reports that in *Hydrolagus* fibres of this muscle insert into the mandibular adductor muscles – we can find no evidence for this in *Callorhinchus*.

***M. labialis anterior*** (Fig. 4c; *m. lab. ant.*)

*Description:* A small, thin muscle with an origin on the posterior side of the prelabial cartilage, ventrally to the insertion of the *m. levator anguli oris anterior pars rostralis.* It inserts on the dorsal process of and along the dorsal side of the superior maxillary cartilage.

*Innervation*: Trigeminal (V) nerve (Didier, 1995).

*Remarks:* Didier (1995) reports that the origin of this muscle is instead on the anterior tip of the prelabial cartilage. This is plausibly consistent with our data as the origin could feasibly wrap around the element in thin connective tissue, but this not visible due to the resolution of our scan.

***M. intermandibularis*** (Figs. 4c, 5b; *m. intermand.*)

**Figure 5.**
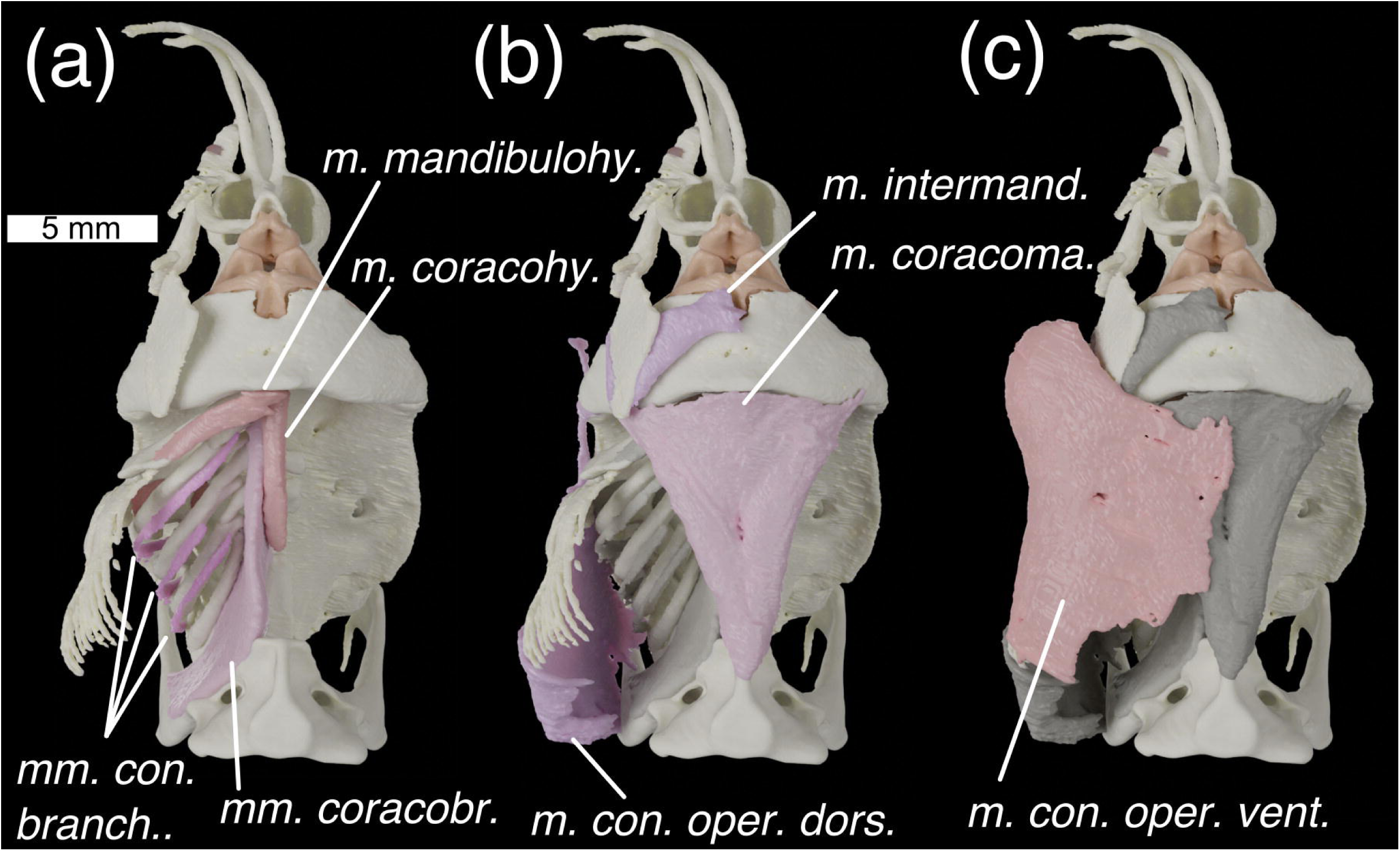
Ventral view of the head of *Callorhinchus milii* with (a) deepest muscles, (b) deeper muscles, and (c) shallow muscles overlain. Colours and abbreviations as in Fig. 4.

*Description:* A thin muscle with an origin along the posterior end and on the dorsal process of the superior maxillary cartilage. It travels ventrally, and posteriorly to the inferior maxillary cartilage has a marked posterior kink, corresponding to an internal sheet of tissue separating the muscle into dorsal and ventral parts. It then travels posterior-ventrally where it is interrupted by an insertion on the posterior and medial edges of the premandibular cartilage. It then continues anteriorly to insert on the mandibular symphysis.

*Innervation:* Trigeminal (V) nerve (Didier, 1995).

*Remarks:* This muscle has been variously reported as two separate muscles (Edgeworth, 1902; Kesteven, 1933; Luther, 1909a), or a single muscle interrupted at the premandibular cartilage (Didier, 1995). Here we follow Didier, who puts forward plausible arguments that both of these parts comprise the same muscle. At the premandibular cartilage where the two parts meet, there is no clear distinction between them in our scan data.

***M. superficialis*** (Fig. 4d; *m. superfic.*)

*Description:* This is a large, very thin muscle, the exact boundaries of which are extremely difficult to make out in the scan data. However, it has a preorbital origin on connective tissue overlying the ethmoid crest. It then travels ventrally to insert along the upper lip, superficially relative to the labial cartilages and all other mandibular muscles, and probably also in the rostrum (Didier, 1995).

*Innervation:* Trigeminal (V) nerve (Didier, 1995).

*Remarks:* Didier (1995) reports that this muscle is unique to *Callorhinchus* among chimaeroids.

***M. constrictor operculi dorsalis*** (Fig. 4d, 5b; *m. con. oper. dors*.)

*Description*: A large, superficial muscle forming the opercular flap dorsal to the opercular cartilage. Its origin is at the base of the scapular process, as well as in connective tissue which connects with the notochord (see Didier, 1995) but which is difficult to resolve in our scan data. It inserts on the rim of the operculum, in a mass of connective tissue above the opercular cartilage. The ***M. constrictor operculi dorsalis anterior*** (Fig. 4d, 6c; *m. con. oper. dors*. *ant.*) is a portion of this muscle with an origin on the ventral rim of the orbit. It extends anteriorly via a tendon which splits to insert on the jaw joint and in the connective tissue of the cheek. The boundary between these two muscles is unresolvable in our scan data.

*Innervation:* Facial (VII) nerve (Didier, 1995).

*Remarks:* The *m. constrictor operculi dorsalis anterior* is only present in *Callorhinchus* amongst chimaeroids (Didier, 1995).

***M. constrictor operculi ventralis*** (Figs. 4d, 5c, 6c; *m. con. oper. vent.*)

**Figure 6.**
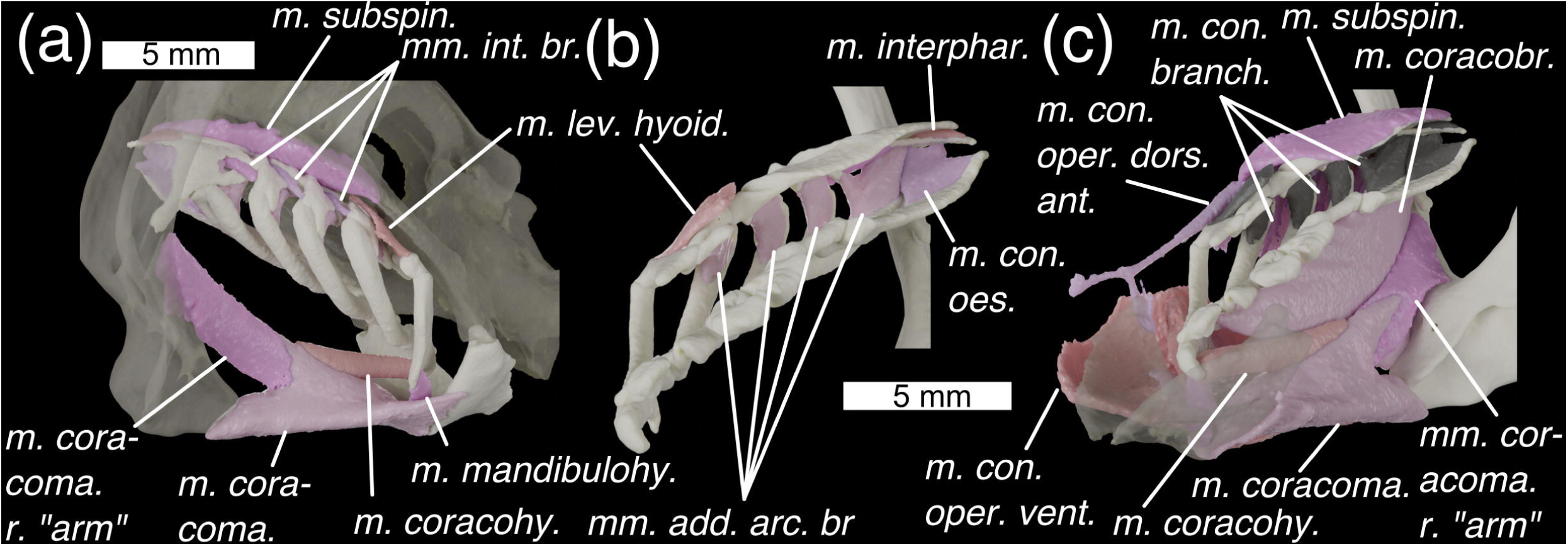
Branchial skeleton of *Callorhinchus milii* with (a) in postero-lateral view with semi-transparent neurocranium and scapulocoracoid, (b) in medial view, and (c) in antero-medial view with a semi-transparent mandible. Colours and abbreviations as in Fig. 3. Additional abbreviations: *m. con. oes., m. constrictor oesophagi; m. coracoma.* r. arm, right “arm” of *m. coracomandibularis* (left not pictured); *m. interphar; m. interpharyngobranchialis*. Otherwise as for Fig. 4.

*Description:* A large superficial muscle in the ventral part of the opercular flap. Its origin is on the opercular cover, ventrally to the opercular cartilage. It wraps around the bottom of the head to insert ventrally to a fascia that joins it to its antimere. A muscle sheet also travels anteriorly, inserting in the connective tissues of the cheek, overlying the mandible and premandibular cartilage. A mesial sheet derived from the muscle travels dorsally to insert in the connective tissue of the *m. mandibulohyoideus,* posteriorly to Meckel’s cartilage.

*Innervation:* Facial (VII) nerve (Didier, 1995).

*Remarks:* Like Didier (1995) we cannot find boundaries between the various component sheets of this muscle described by Kesteven (1933) and consider it to be a single muscle.

***M. levator hyoideus*** (Figs. 4a, 6a,b; *m. lev. hyoid.*)

*Description*: This is a small, thin sheet of muscle, with an origin on the underside of the neurocranium, between the foraminae for the *nervus hyoideo-mandibularis facialis* and the *nervus palatinus facialis*. It inserts along the ventral part of the epihyal’s postero-lateral edge. It is also linked by connective tissue to the opercular cartilage.

*Innervation:* Facial (VII) nerve (Didier, 1995)

***M. mandibulohyoideus*** (Fig. 4a, 5a, 6a; *m. mandibulohy.*)

*Description:* This is a thin muscle with an origin on the posterior symphysis of the mandible, where it meets its antimere as well as the ventral constrictors via a tendon. It inserts on the ventral angle of the ceratohyal.

*Innervation:* The facial (VII) nerve (Didier, 1995) as well as the glossopharyngeal (IX) nerve in *Hydrolagus* (Anderson, 2008).

*Remarks:* Didier (1995) suggests that this muscle is correlated with the evolution of autostyly. Kesteven (1933) called this muscle the geniohyoideus, while Didier (1995) called it the interhyoideus. Anderson (2008) coined the name mandibulohyoideus for it to reflect the muscle’s apomorphy with respect to the interhyoideus and geniohyoideus in osteichthyans, and this is the name we use here.

***Mm constrictores branchiales.*** (Fig. 4b, 5a, 6c; *mm. con. branch.*)

*Description:* We find three branchial constrictor muscles. These are so small and thin as to be difficult to characterise, but their origins are high up on the lateral sides of epibranchials II and III, and on the lateral side of the posterior pharyngobranchial complex. Each extends antero-posteriorly onto the ventral side of the ceratobranchial of the anterior arch (i.e. ceratobranchials I-III).

*Innervation:* Glossopharyngeal (IX) and vagus (X) nerves (Didier, 1995).

*Remarks:* Like Didier (1995) we were unable to find the fourth branchial constrictor that Edgeworth (1935) described, however, the constrictor muscles are so thin that it is possible that it is unresolved in the scan data. Like Edgeworth (1935) reported, each constrictor passes between two arches. We agree with Didier that the ventral lengths of these muscles are likely described by Kesteven (1933) as possible *transversi ventrales*: “three long slender muscles, each of which arises from each of the first three basi-branchial cartilages and extends along the outer curve of the ceratobranchial of the same arch.”. Kesteven (1933) also describes three sets of “dorsal oblique interarcual muscles”. Of these the “external dorsal oblique muscles” seem likely to be the dorsal part of the branchial constrictors on the basis of descriptions as a “short muscle which arises from the dorsal surface of the epibranchial of the fourth, third, and second arches, and is inserted on the top of the cerato-branchial of the third, second, and third arches”.

***Mm. interarcuales branchiales*** (Fig. 4a, 6a; *mm. int. br.*)

*Description:* These muscles are very small and extend from the ventral surface of the pharyngobranchials to the posterior edge of the epibranchials I-III.

*Innervation:* Glossopharyngeal (IX) and vagus (X) nerves (Didier, 1995; Edgeworth, 1935)

*Remarks:* These muscles are not mentioned by Didier (1995) but appear to roughly match the description of the external dorsal oblique muscles by Kesteven (1933), with the difference that they travel from pharyngobranchial to epibranchial rather than from epibranchial to ceratobranchial.

***Mm adductores arcuum branchiales*** (Figs. 4a, 6b; *mm. add. arc. br.*)

*Description:* These muscles lie medial to the first four branchial arches, extending between the dorsal surface of the ceratobranchials and the ventral surface of the epibranchials. The origin of the first three is on epibranchials I-III, and they insert on ceratobranchials I-III. The fourth has two origins on the posterior pharyngobranchial complex (pharyngobranchials IV-V) and inserts on ceratobranchials IV-V. Also associated with the fourth is the ***M. constrictor oesophagi*** (Fig. 6b; *m. con. oes.*), the origin of which lies along the posterior process of the posteriormost pharyngobranchial, and which inserts along the postero-lateral length of the basibranchial copula.

*Innervation:* Glossopharyngeal (IX) and vagus (X) nerves (Didier, 1995; Edgeworth, 1935).

*Remarks:* These are probably the lateral internal dorsal oblique muscles of Kesteven (1933), although our data shows them to insert high up on the epibranchial rather than on the pharyngobranchials as described by Kesteven.

***M. cucullaris superficialis*** (Fig. 4d; *m. cuc. sup*.)

*Description:* A large muscle with a broad origin on the postorbital crest and overlying the epaxial muscles. It inserts on the scapular process, dorsal to the origin of the *M. constrictor operculi dorsalis*.

*Innervation:* 4th branch of the Vagus (X) nerve (Edgeworth, 1935).

***M. protractor dorsalis pectoralis*** (Fig. 4b; *m. prot. dors. pect.*)

*Description:* This muscle has its origin on the posterior part of the orbital process and inserts on the anterior edge and medial face of the scapular process. Its boundary with the *m. retractor dorsalis pectoralis* is difficult to distinguish in the scan data.

*Innervation:* Glossopharyngeal (IX) and/or the vagus nerve (X) (Ziermann et al., 2014).

*Remarks:* There is some disagreement over whether this muscle is a trunk muscle or a branchial muscle (see Ziermann et al., 2014), which arises from uncertainty over the innervation.

***M. cucullaris profundus*** (Fig. 4c; *m. cuc. prof.*) is a short, thin muscle. It has its origin on the underside of the otic region, lateral to the *M. subspinalis*. It inserts on the postero-ventral end of the posterior pharyngobranchial complex’s lateral side, latero-ventral to the ceratobranchials.

*Innervation:* 3rd branch of the vagus (X) nerve (Edgeworth, 1935).

***M. subspinalis*** (Fig. 4b, 6a,c; *m. subspin.*)

*Description:* Broad and flat with an origin along a stretch of the otic shelf, medial to that of the *m. cucullaris profundus*. It inserts along the dorsal surfaces of pharyngobranchials I and II.

*Innervation:* Spinal nerves, specifically the *plexus cervicalis*, formed by two or more anterior spinal nerves (Edgeworth, 1935).

***M. interpharyngobranchialis*** (Fig. 6b; *m. interphar.*)

*Description:* This is a very small muscle that joins the second pharyngobranchial to the posterior pharyngobranchial complex.

*Innervation:* Spinal nerves, specifically the *plexus cervicalis*, formed by two or more anterior spinal nerves (Edgeworth, 1935).

*Remarks:* Edgeworth (1935) describes this muscle, while Didier (1995) describes it as absent. It is of such a small size that it might be easily missed.

***M. coracomandibularis*** (Figs. 4a, 5b, 6a,c; *m. coracoma.*)

*Description:* A very large muscle with its main origin on the T-shaped antero-ventral face of the coracoid region of the pectoral girdle, and which inserts along Meckel’s cartilage. Viewed ventrally the muscles is triangular, broadening anteriorly. As described by Shann (1919) and Didier (1995) the muscle is split into shallow and deep portions – the deeper portion inserts along the posterior of the mandible as a sheet, while the shallower portion splits into two thick bundles, which diverge about halfway along the muscles length to insert at either side of Meckel’s cartilage. At its anterior extent the muscle is fairly flat, but towards the origin it develops a pronounced dorsal keel oriented postero-dorsally. This keel is cleft by a v-shaped septum, where the muscle splits into paired “arms” (Fig. 6a,c) that diverge laterally to origins on the left and right bases of the scapular processes, dorsal to the pectoral fin articulations.

*Innervation:* Spinal nerves, specifically the *plexus cervicalis*, formed from two or more anterior spinal nerves (Edgeworth, 1935).

*Remarks:* The morphology of this muscle varies in different holocephalan genera (Didier, 1995; Shann, 1919), particularly its relationship to the pectoral symphysis. Both Shann and Didier describe a v-shaped septum in the *M. coracomandibularis* of *Chimaera*, but report that this cannot be found in *Callorhinchus.* Our scan data shows it to be present, illustrated in our figures by the junction between the two colours of the muscle. As in Shann s description of *Chimaera,* the spinal nerves that innervate the *M. coracomandibularis* enter the muscle at this septum.

***M. coracohyoideus*** (Figs. 4a, 5a,6a,c; *m. coracohy.*)

*Description:* A long thin muscle with its origin on the dorsal surface of the *M. coracomandibularis,* anteriorly to the V-shaped septum. It then extends anteriorly to insert on the posterior side of the basihyal.

*Innervation:* Spinal nerves, specifically the *plexus cervicalis*, formed by two or more anterior spinal nerves (Edgeworth, 1935)

*Remarks:* Didier (1995) notes that it is unclear whether this muscle takes origin from the coracoid or *m. coracomandibularis* in *Callorhinchus*. Our scan data shows that its entire origin lies on the *m. coracomandibularis* (Fig. 4a).

***Mm. coracobranchiales*** (Fig. 4b, 5a, 6c; *mm. coracobr.*)

*Description:* Long, thin muscles, with an origin along the dorso-lateral corner of the coracoid and the base of the scapular process. They insert ventrally on the hypobranchials. The anterior three attach to hypobranchials I-III, while the fourth and fifth attach to the fourth, posteriormost, hypobranchial.

*Innervation:* Spinal nerves, specifically the *plexus cervicalis*, formed by two or more anterior spinal nerves (Edgeworth, 1935).

*Remarks:* Although the muscles have separate heads, they are difficult to separate in our scan data and have been segmented out together.

***M. epaxialis*** (Fig. 4d, *m. epaxialis*)

*Description:* Sheet-like muscle with origin on the top of the head, along the dorsal ridge and above the orbit. It inserts posteriorly with the dorsal myomeres.

*Innervation:* Spinal nerves (Edgeworth, 1935).

*Remarks:* Unlike in *Scyliorhinus* where the epaxials terminate posterior to the orbit, in *Callorhinchus* they extend well anteriorly, terminating in front of the orbits.

***M. retractor dorsalis pectoralis*** (Fig. 4b, *m. ret. dors. pect.*)

*Description:* This has its origin on the posterior dorsal part of the scapulocoracoid, and along the bottom of the filament which extends posteriorly from the dorsal tip of the scapular process. It extends posteriorly to insert in the trunk musculature.

*Innervation:* Spinal nerves (Didier, 1995).

***M. retractor latero-ventralis pectoralis*** (Fig. 4b,c, *m. ret. lat.-vent. pect. l+m*)

*Description:* This muscle comprises two parts. These have their origin laterally and medially on the scapular process. They insert posteriorly, in the dorsal muscle tissue of the body cavity.

*Innervation:* Glossopharyngeal (IX) and/or the vagus nerve (X) (Ziermann et al., 2014).

*Remarks:* There is some disagreement over whether this muscle is a trunk muscle or a branchial muscle (see Ziermann et al. (2014), which arises from uncertainty over the innervation.

***M. retractor mesio-ventralis pectoralis*** (Fig. 4c, *m. ret. mes.-vent. pect.*)

*Description:* This is a sheet of muscle. Its origin is in several places at the bottom of the scapular process and on the posterior of the coracoid. It inserts in the lateral and ventral muscle of the body.

*Innervation:* Glossopharyngeal (IX) and/or the vagus nerve (X) (Ziermann et al., 2014).

*Remarks:* As above here is some disagreement over whether this muscle is a trunk muscle or a branchial muscle (see Ziermann et al. (2014)).

***M. rectus dorsalis*** (Fig. 3c; *m. rect. dors.*)

*Description:* This is antagonistic to the *M. rectus ventralis*. It has its origin in the orbit anteriorly to the trigeminal nerve entrance. It inserts dorsally around the eye.

*Innervation:* Oculomotor (III) nerve (Edgeworth, 1935).

***M. rectus ventralis*** (Fig. 3c; *m. rect. vent*.)

*Description:* This muscle is antagonistic to the *M. rectus dorsalis*. It has its origin in the orbit anteriorly to the trigeminal nerve entrance, ventrally the profundus (V) nerve. It inserts ventrally around the eye, with a comparatively broad insertion.

*Innervation:* Oculomotor (III) nerve (Edgeworth, 1935).

***M. rectus lateralis*** (Fig. 3c; *m. rect. lat*.)

*Description:* This muscle is antagonistic to the *M. rectus medialis.* has its origin in the orbit anteriorly to the trigeminal nerve entrance. It inserts posteriorly around the eye.

*Innervation:* Abducens (VI) nerve (Edgeworth, 1935).

***M. rectus medialis*** (Fig. 3c; *m. rect. med*.)

*Description:* This muscle has its origin relatively anteriorly to the other rectus muscles, and is antagonistic to the *M. rectus lateralis.* It inserts anteriorly around the eye.

*Innervation:* Oculomotor (III) nerve (Edgeworth, 1935).

***M. obliquus ventralis*** (Fig. 3c; *m. obl. vent.*)

*Description:* This muscle is antagonistic to the *M. obliquus dorsalis*. It has its origin just ventral to the antorbital crest, well anteriorly in the orbit. It inserts antero-ventrally around the eye.

*Innervation:* Oculomotor (III) nerve (Edgeworth, 1935).

***M. obliquus dorsalis*** (Fig. 3c; *m. obl. dors.*)

*Description:* This muscle is antagonistic to the *M. obliquus ventralis*. It has its origin just medial to the antorbital crest, by the ophthalmic foramen. It inserts antero-dorsally around the eye.

*Innervation:* Trochlear (IV) nerve (Edgeworth, 1935).

#### Ligaments

Didier identifies two paired ligaments in the snout of *Callorhinchus*:

***Ligamentum labialis*** (Fig. 2a; *lig. lab.*)

*Description:* This travels between a dorsal process on the nasal capsule and then wraps around the antero-ventral part of the prelabial cartilage.

*Remarks:* As Didier (1995) notes this is only present in *Callorhinchus*.

***Ligamentum rostralis*** (Fig. 2a; *lig. rost.*)

*Description:* This travels between the dorso-anterior part of the prelabial cartilage onto the lateral rostral rod.

*Remarks*: Didier (1995) notes this is only present in *Callorhinchus*.

***?Ligamentum pharyngohyoideus*** (Fig. 1d; *?lig. pharh.*)

*Description*: A very small ligament (or possibly a slip of muscle) with an origin on the neurocranial floor, posterior to the quadrate processes and lateral to the hypophyseal notch, and inserting on the pharyngohyal dorsally.

*Innervation*: This is presumably innervated by the facial (VII) nerve, given its location, but no precise innervation can be established.

*Remarks*: This structure has not been previously reported. Given that it has not been observed before, and its poor visibility in the dataset, we are cautious its definite existence pending its discovery in gross dissection. However, it is present on both sides of the dataset. Based on the apparent lack of fibres we presume this is a ligament rather than a muscle, but the latter is certainly not impossible. It may play a role in the operation of the operculum.

### Scyliorhinus canicula

#### Cranial Cartilages

The cranial skeleton of *Scyliorhinus canicula* comprises the neurocranium (Fig. 7), palatoquadrates, Meckel’s cartilages, two pairs of labial cartilages, the hyoid arch, and five branchial arches (Fig. 8). As in other elasmobranchs the branchial skeleton stretches well-posterior to the neurocranium, bounded posteriorly by ventrally joined scapulocoracoids.

**Figure 7.**
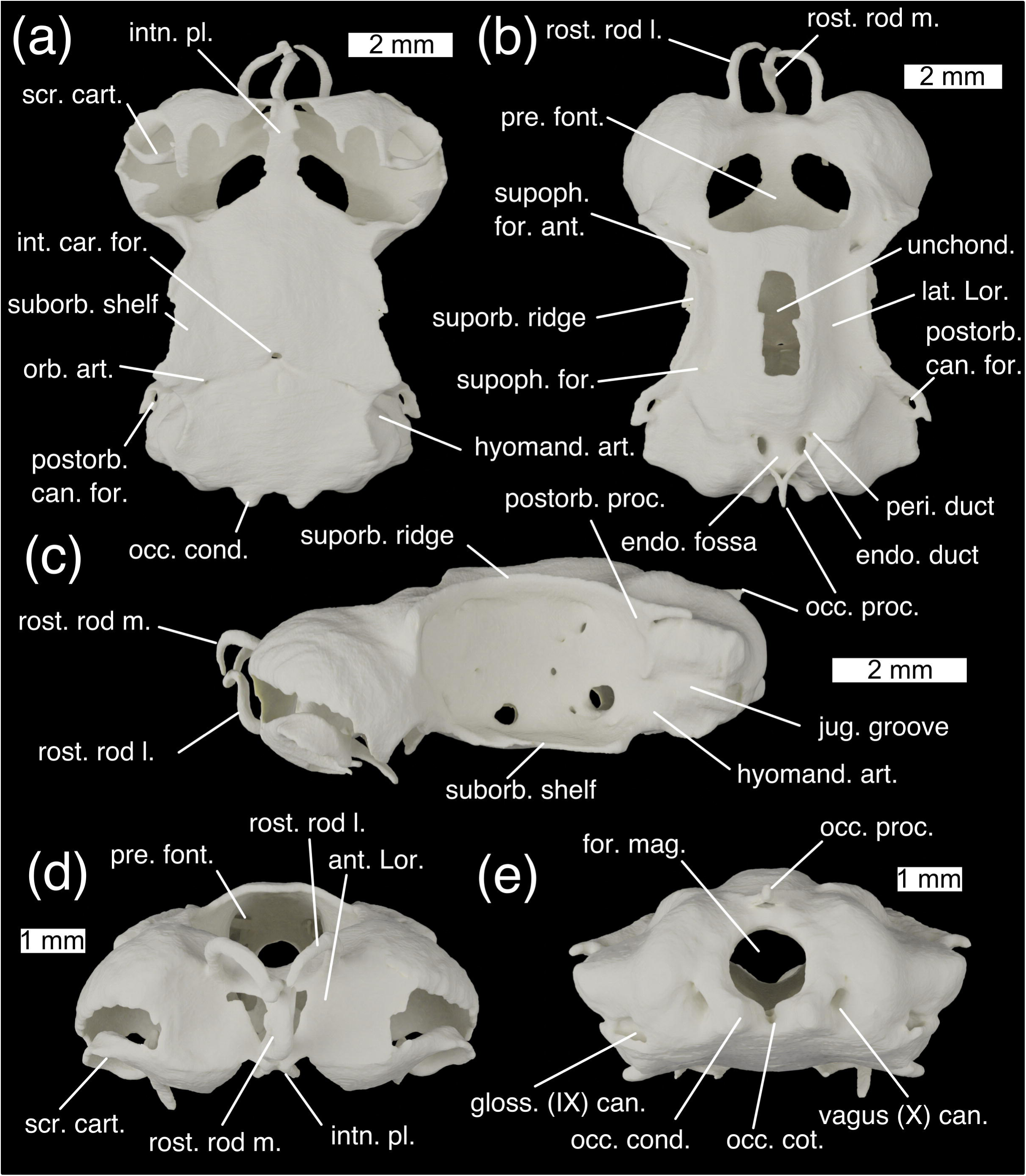
The neurocranium of *Scyliorhinus canicula* in (a) ventral, (b) dorsal, (c) lateral, (d) anterior, and (e) posterior views. Abbreviations: ant. Lor., anterior depression for ampullae of Lorenzini; endo. duct, endolymphatic duct; endo. fossa, endolymphatic fossa; for. mag., foramen magnum; gloss. (IX) can., glossopharyngeal (IX) nerve canal exit; hyomand. art, hyomandibular articulation surface.; int. car. for., foramen for the internal carotids; intn. pl., internasal plate; jug. groove, jugular groove; lat. Lor., lateral furrows for ampullae of Lorenzini; postorb. can. for., foramen for postorbital sensory canal; occ. cond., occipital condyle; occ. cot., occipital cotylus; occ. proc., occipital processes; orb. art., foramen for the orbital artery; peri. duct, perilymphatic duct; postorb. can. for., foramen for postorbital sensory canal; postorb. proc., postorbital process; pre. font., precerebral fontanelle; rost. rod l, lateral rostral rod; rost. rod m., median rostral rod; scr. cart., scrolled cartilage; suborb. shelf;supoph. for. foraminae for twigs of the superophthalmic complex; supoph. for., foraminae n for the superophthalmic complex; supoph. for. ant., anterior foramen for the superophthalmic complex; suporb. ridge, supraorbital ridge; unchond., incompletely chondrified area; vagus (X) can., vagus (X) nerve canal exit

**Figure 8.**
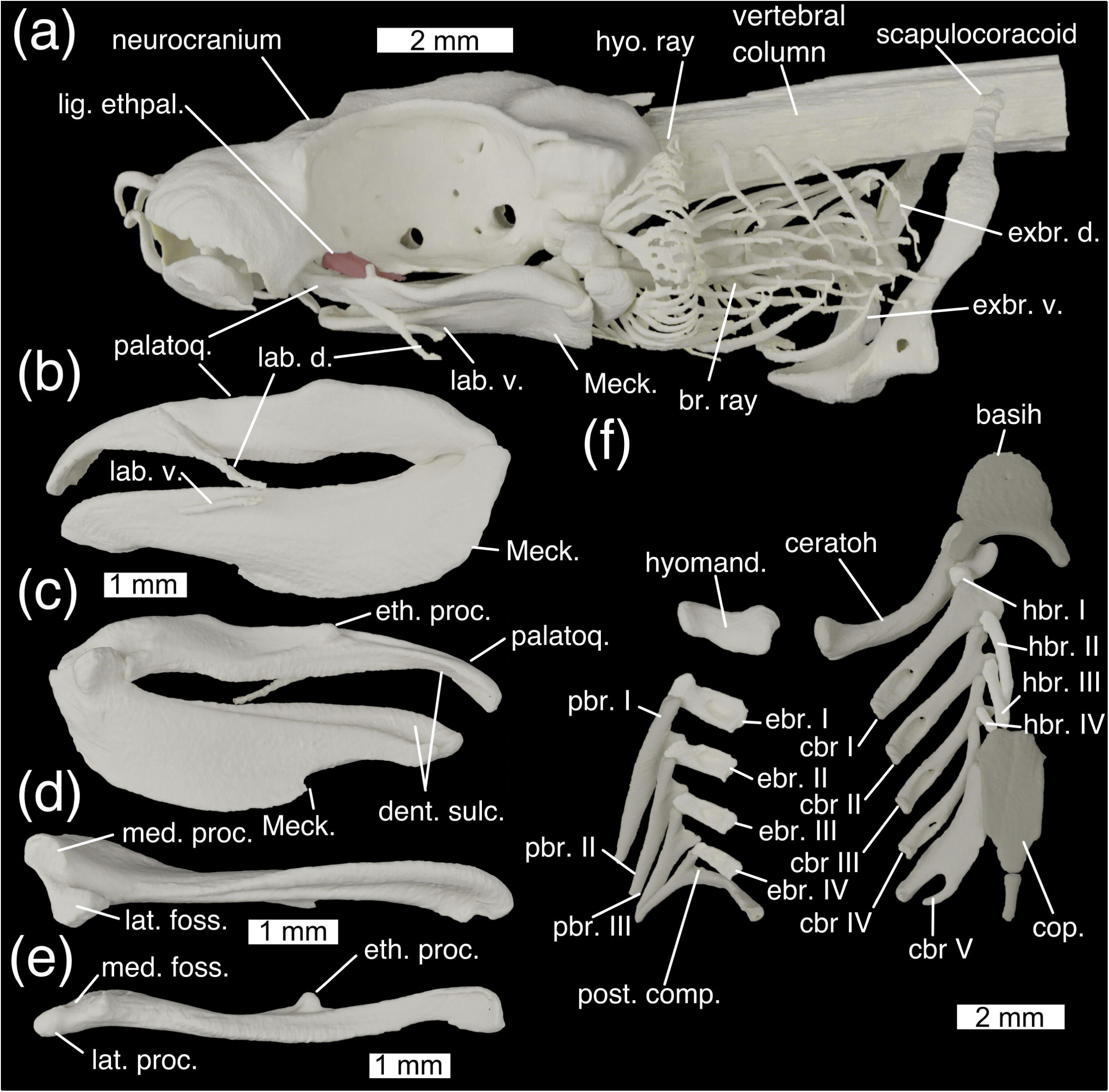
Cranial skeleton of *Scyliorhinus canicula* with (a) whole cranial skeleton in lateral view, mandibles in (b) lateral and (c) medial view, (d) left Meckel’s cartilage in dorsal view, (e) left palatoquadrate in ventral view, (f) dorsal gill skeleton in ventral view and (g) ventral gill skeleton in dorsal view. Abbreviations: basih., basihyal; br. ray; branchial rays; ceratoh, ceratohyal; cbr., ceratobranchial; cop., basibranchial copula; dent. sulc., dental sulcus; lab. d., dorsal labial cartilage; eth. proc., ethmoid process; ebr, epibranchial; exbr. d., dorsal extrabranchial cartilages; exbr. v., ventral extrabranchial cartilages; hyomand., hyomandibula; hyo. ray, hyoid rays; hbr., hypobranchial; lab. v., ventral labial cartilage; lat. foss., lateral fossa of Meckel’s cartilage; lat. proc., lateral process of palatoquadrate; Meck., Meckel’s cartilage; med. foss, medial fossa of palatoquadrate; med. proc., medial process of Meckel’s cartilage; palatoq., palatoquadrate; pbr., pharyngobranchial; post. comp., posterior epibranchial/pharyngobranchial complex

The **neurocranium** in *Scyliorhinus* is fairly flat, with large orbits and broad ethmoid and otic regions. The olfactory capsules are subspherical and very large, taking up about a third of the volume of the entire neurocranium. Ventrally they are open and partially covered by digitate and scrolled projections of cartilage (Fig. 7a, scr. cart.). They are separated by a medial wall of cartilage which expands ventrally into a narrow internasal plate (Fig. 7a, intn. pl.). Anteriorly the olfactory capsules are marked by shallow depressions, in which an anterior grouping of the ampullae of Lorenzini sits (Fig. 7d, ant. Lor.). This structure is supported by three rostral rods (Fig. 7b, rost. rod m., l.), one unpaired ventrally and one paired dorsally, which curve, converging centrally. Between the two olfactory cartilages dorsally a large precerebral fontanelle is situated (Fig. 7b, d, pre. font.).

The orbits are large and oval, comprising about half of the length of the neurocranium. Dorsally they are bounded by a strong supraorbital ridge (Fig. 7b, suporb. ridge), with sharp anterior and posterior terminations. Posteriorly this forms the postorbital process (Fig. 7c, postorb. proc.) which, like in other elasmobranchs, does not extend ventrally to form a postorbital arcade. Between the orbits the roof of the neurocranium rises to form a shallow ridge, the apex of which is incompletely chondrified (Fig. 7b, unchond.). Between this and the supraorbital ridges are a pair of shallow furrows carrying ampullae of Lorenzini (Fig. 7b, lat. Lor.), which are innervated by twigs of the superficial ophthalmic complex through foraminae in the supraorbital ridge (Fig. 7b., supoph. for.). The main trunk of the superficial ophthalmic complex enters the orbit through a large foramen postero-dorsally (Fig. 9a; supoph. for. post.). A small foramen next to this permits the entry of the profundus into the orbit (Fig. 9a; prof. for.). The superficial ophthalmic complex and profundus exit the orbit together passing through a large foramen in the dorso-anterior corner onto the anterior neurocranial roof (Figs. 7b, 9a; supoph. for. ant.). Posterior to this, high up on the wall of the orbit is a small foramen through which the abducens (IV) nerve enters the orbit (Fig. 9a; abd. (IV) for.) (Holmgren, 1940). Ventrally the orbit is bounded by a broad suborbital shelf (Fig. 7a, suborb. shelf). In the postero-ventral corner of the orbit is a large foramen through which the facial (VII) and trigeminal (V) nerves enter the orbit (Fig. 9a; V+VII). Ventro-laterally to this is a small foramen through which the orbital artery enters the orbit (Fig. 9a; orb. art.). Antero-dorsally to the facial and trigeminal nerve foramen is an opening for the oculomotor (III) nerve (Fig. 9a; oculom. (III) for.) (Holmgren, 1940), while anteriorly to it is a foramen for the pituitary vein (Fig. 9a; pit. v.) (Holmgren, 1940). Anteriorly to this is a foramen for the efferent pseudobranchial artery (Fig. 9a; eff. pseud.), and anteriorly to this the foramen for the optic (II) nerve (Fig. 9a; opt. (II) for.) as well as the optic artery. A small foramen antero-dorsal to the foramen for the optic nerve permits entry for the anterior cerebral vein (Fig. 9a; ant. cer. v.). A foramen in the antero-ventral corner of the orbit provides an entry for the nasal vein through the orbitonasal canal (Fig. 9a; orbnas. can.). Posteriorly to the orbit on the skull roof is a large foramen, through which the postorbital sensory canal passes (Fig. 7, post. can. for.).

**Figure 9.**
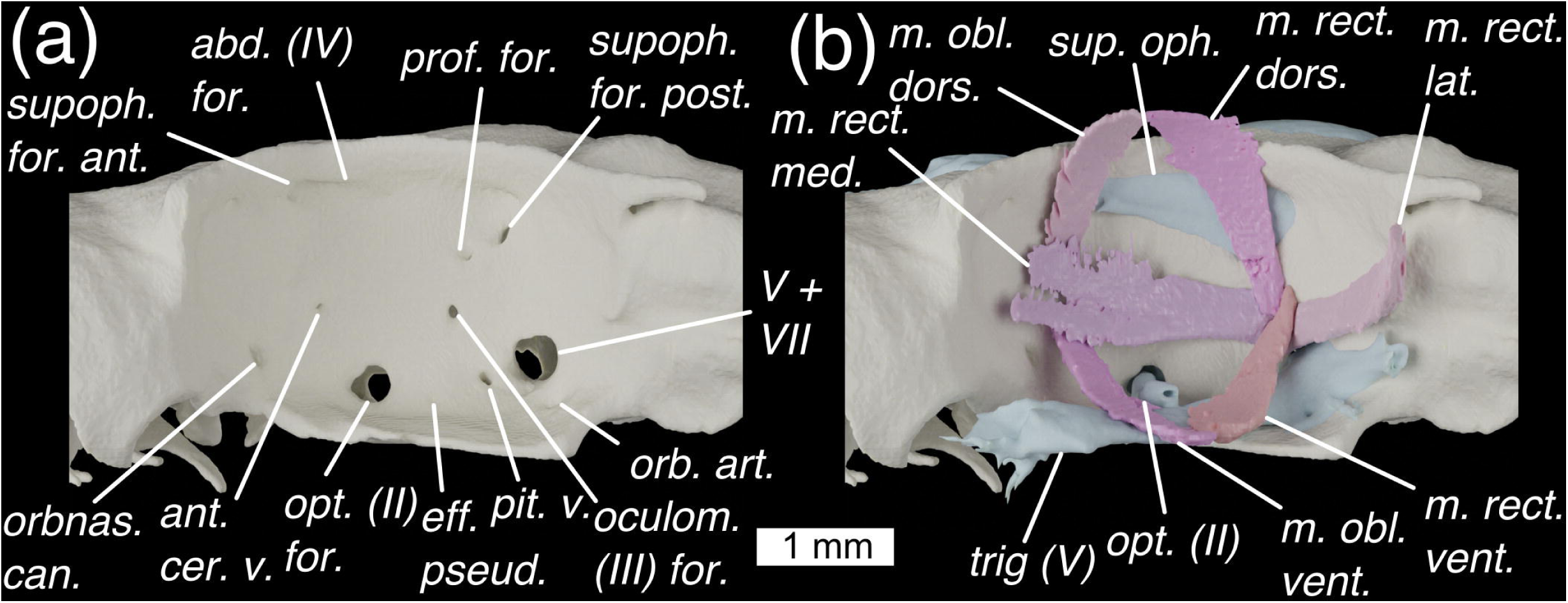
The orbit of *Scyliorhinus canicula* shown in lateral view with (a) foraminae shown, and (b) external optic muscles and cranial nerves. Colours as in Fig. 9 with light-blue for cranial nerves. Abbreviations: abd (IV) for., foramen for the abducens (IV) nerve; ant. cer. v., foramen for the anterior cerebral vein; eff. pseud, foramen for the efferent pseudobranchial artery; *m. obl. dors.*, *m. obliquus dorsalis*; *m. obl. vent.*, *m. obliquus ventralis*; *m. rect. dors.*, *m. rectus dorsalis*; *m. rect. lat.*, *m. rectus lateralis*; *m. rect. med.*, *m. rectus medialis*; *m. rect. vent.*, *m. rectus ventralis*; oculom. (III) for., foramen for the oculomotor (III) nerve; opt (II), optic nerve; orbnas. can., foramen for the orbitonasal canal; opt. (II) for., foramen for the optic (II) nerve; orb. art., foramen for the orbital artery; pit. v., foramen for pituitary vein; prof. for., foramen for profundus; supoph. for. ant., anterior foramen for the superophthalmic complex; sup. oph., superficial ophthalmic complex (V); supoph. for. post., posterior foramen for the superophthalmic complex; V+VII, foramen for entry of trigeminal (V) and facial (VII) nerves into the orbit; trig. (V), trigeminal nerve

The otic capsules are broad and marked by a dorsal ridge formed by the anterior and posterior semicircular canals, with the external semicircular canal forming a pronounced lateral ridge. Below the lateral ridge is a marked groove for the jugular vein (Fig. 7c, jug. groove). Anteroventrally to this lies a flat surface on which the hyomandibula articulates (Fig. 7c, hyomand. art.). Posteriorly to this jugular groove is the exit point of the glossopharyngeal (IX) nerve canal (Fig. 7e, gloss. (IX) can.). Between the dorsal ridges is a shallow endolymphatic fossa (Fig. 7b, endo. fossa), containing paired openings – a larger pair for the endolymphatic ducts (Fig. 7b, endo. duct) and a smaller pair antero-laterally for the perilymphatic ducts (Fig. 7b, peri. duct). Posteriorly to these are paired occipital processes that join to form an arc (Fig. 7b, c, occ. proc.). A rounded foramen magnum (Fig. 7e, for. mag.) is positioned below this, and ventrally to this is a shallow occipital cotylus (Fig. 7e, occ. cot.), bounded laterally by rounded occipital condyles (Fig. 7e, occ. cond.). Laterally to these are a pair of foraminae for the exit of the Vagus (X) nerve canal (Fig. 7e, vagus (X) can.). At least one of the spinal nerves appears to join the Vagus to leave through the vagal canal: the rest diverge posteriorly to the braincase.

The ventral side of the neurocranium is flat, broad, and fairly featureless. At about one third of the length from the posterior it is punctured by a medial foramen through which the internal carotids enter the neurocranium (Fig. 7a, int. car. for.), and lateral to this are paired foraminae through which the orbital arteries enter the orbits (Fig. 7a; orb. art.).

*Scyliorhinus* also possesses a small prespiracular cartilage, in the anterior wall of the spiracle (Ridewood, 1896, Tomita *et al*. 2018). However, resolution surrounding the spiracle in our dataset proved insufficient to locate this.

The **palatoquadrates** (Fig. 8a, b; palatoq.) are about one third of the length of the neurocranium and joined at an anterior symphysis. They are low and flat, with a short, rounded ethmoid process on the medial face of the palatine process, via which the neurocranium is joined to the palatoquadrate by the ethmopalatine ligament (see below) (Fig. 8e; eth. proc.). Anteriorly to this process the dorsal edge is marked by a shallow groove. The inside edge carries a shallow sulcus for the teeth (Fig. 8c; dent. sulc.). **Meckel’s cartilages** are about one and a half times as deep as the palatoquadrates (Fig. 8a, b; Meck.), and are joined at an anterior symphysis. Dorsally it is grooved by a sulcus for the attachment of teeth (Fig. 8c; dent. sulc.). It is ventrally tall, particularly along its posterior half before abruptly losing height. A dorsal and a ventral pair of labial cartilages are positioned lateral to the jaws, which together form a v-shape with the open end anteriorly (Fig. 8a, b; lab. d., lab. v.). The elements have a double-articulation. A large medial process (Fig. 8d; med. proc.) and lateral fossa (Fig. 8d; lat. foss.) on Meckel’s cartilage articulate with a narrow lateral process (Fig. 8e; lat. proc.) and shallow medial fossa (Fig. 8e; med. foss.) on the palatoquadrate.

The hyoid arch comprises a basihyal, and paired ceratohyals and hyomandibulae (epihyals). The basihyal is broad and flat (Fig. 8f; basihy.), and is punctured centrally by a single foramen for the thyroid gland stalk (De Beer and Moy-Thomas, 1935). A rim curves around its anterolateral edge, and terminates posteriorly, forming paired ceratohyal articulations along with posteriorly projecting paired processes. The ceratohyal (Fig. 8f; ceratoh.) is laterally flattened and curved dorsally. The anterior end is expanded into two heads, the anterior of which articulates in the basihyal’s fossa. The hyomandibula (Fig. 8f; hyomand.) is short and stout, with expanded ends for the articulation with the braincase and the ceratohyal. It articulates on the ventral side of the otic capsule, immediately posterior to the orbit. Hyoid rays (Fig. 8f; hyo. ray) are attached to the posterior side of the hyomandibula and ceratohyal, and form a branching series of rays that support the first gill flap.

Posterior to the hyoid arch are five branchial arches. The floor of the pharynx is supported by a basibranchial copula and four hypobranchials. The basibranchial copula (Fig. 8f; cop.) is a large, flat, posteriorly located element with a posterior tail, the posterior length of which is mineralised. The anteriormost hypobranchial (Fig. 8f; hbr.) is small and cuboid, and oriented anteriorly, overlying the ceratohyal and joining the posterior process of the basihyal to the first ceratobranchial. The posterior three hypobranchials are long and thin, each smaller than the one before, and are oriented posteriorly towards the anterior edge of the copula from the junction between the first and second, second and third, and third and fourth ceratobranchials. The first four ceratobranchials (Fig. 8f; cbr.) are long and thin, and their distal end is expanded into two heads, each of which meets the hypobranchial and ceratobranchial of the arches in front and behind. The dorsal proximal surface is marked by a deep spoon-shaped concavity in which the branchial adductor muscles sit, and which is pierced by a foramen, possibly to allow vascularization or innervation of the adductor muscles. The posteriormost ceratobranchial is broad and flat, lacks a distal head, and has its proximal end branched into two parts. The first four epibranchials (Fig. 8f; ebr.) are short and rectangular with a short anterior process, and concave ventrally for the branchial adductors. They are pierced by a foramen, again possibly for the vascularization or innervation of the branchial adductors. There are three separate pharyngobranchials (Fig. 8f; pbr.), which are long, thin arrowheads swept posteriorly. Their proximal ends are expanded in two heads, the anterior one of which articulates with the anterior epibranchial and the posterior one of which overlies the epibranchial behind. The posteriormost pharyngobranchial(s) and the fifth epibranchial are fused into a single pick-shaped posterior complex (Fig. 8f; post. comp.) with an anterior process articulating with the fourth epibranchial, a ventral process articulating with the fifth ceratobranchial, and a posterior swept back process. The branchial rays (Fig. 8a; br. ray) are unbranched, unlike those of the hyoid, and attached to the ceratobranchials and epibranchials of the first four branchial arches, becoming less numerous on posterior arches. Five dorsal extrabranchial cartilages (Fig. 8a; exbr. d.) overlie each gill flap, with heads on the lateral face of the expaxial (I) and cucullaris (II-V) muscle. There are three ventral extrabranchial cartilages (Fig. 8a; exbr. v.), supporting the second, third, and fourth gill flaps ventrally. These have complex-shaped heads that are ventrally inserted between the coracobranchial muscles and over lying the coracoarcual muscles.

#### Cranial Muscles

This account follows the terminology of Edgeworth (1935).

***M. levator labii superioris*** (Fig. 10a; *m. lev. lab. sup.*)

**Figure 10.**
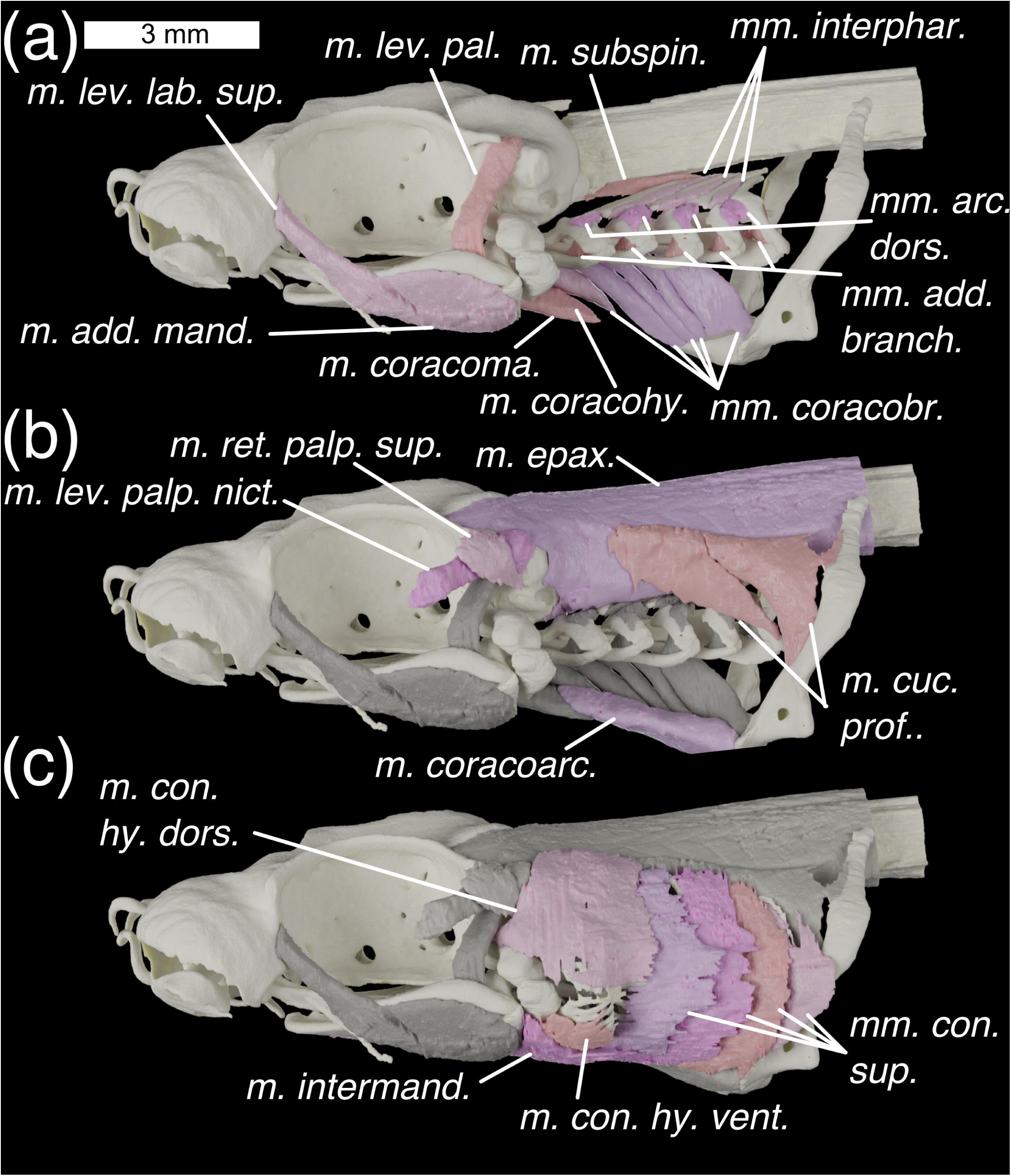
Lateral view of the head of *Scyliorhinus canicula* with (a-c) progressively shallower muscles shown. Colours: cream, cartilage; beige, pinks, muscles; greys, deeper muscles. Abbreviations: *mm. add. branch., m. adductors branchiales; m. add. mand., m. adductor mandibulae*; *mm. arc. dors., mm. arcuales branchiales; m. con. hy. dors. m. constrictor hyoideus dorsalis; m. con. hy. vent. m. constrictor hyoideus ventralis; mm. con. sup., mm. constrictors superficiales; m. coracoarc., m. coracoarcualis; mm. coracobr., mm. coracobranchiales; m. coracohy., m. coracohyoideus; m. coracoma., m. coracomandibularis; m. cuc. prof., m. cucullaris profundus; m. epax., m. epaxialis; mm. interphar., mm. interpharyngobranchiales; m. intermand., m. intermandibularis*; *m. lev. lab. sup., m. levator labii superioris*; *m. lev. pal., m. levator palatoquadrate; m. lev. palp. nict., m. levator palpebrae nictitantis; m. subspin., m. subspinalis; m. ret. palp. sup., m. retractor palpebrae superioris*

*Description:* A long thin muscle with its origin on the posterior part of the nasal capsule, immediately antero-lateral to the orbit. It extends postero-ventrally to insert in the fibres of the dorsal part of the adductor muscle. It is separated from the *M. adductor mandibulae* by the trigeminal (V) nerve, which lies across the lateral surface, but the border between the fibres of the two muscles are largely indistinguishable in our scan data and are segmented as one model. Fibres of the *m. levator labii superioris* reach the level of the fascia separating the dorsal and ventral parts of the mandibular adductor.

*Innervation:* Trigeminal (V) nerve (Edgeworth, 1935).

*Remarks:* This muscle is variable in galeomorph sharks, with phylogenetic significance (See discussion in Soares and Carvalho, 2013).

***M. adductor mandibulae*** (Figs. 10a, 11a; *m. add. mand.*)

**Figure 11.**
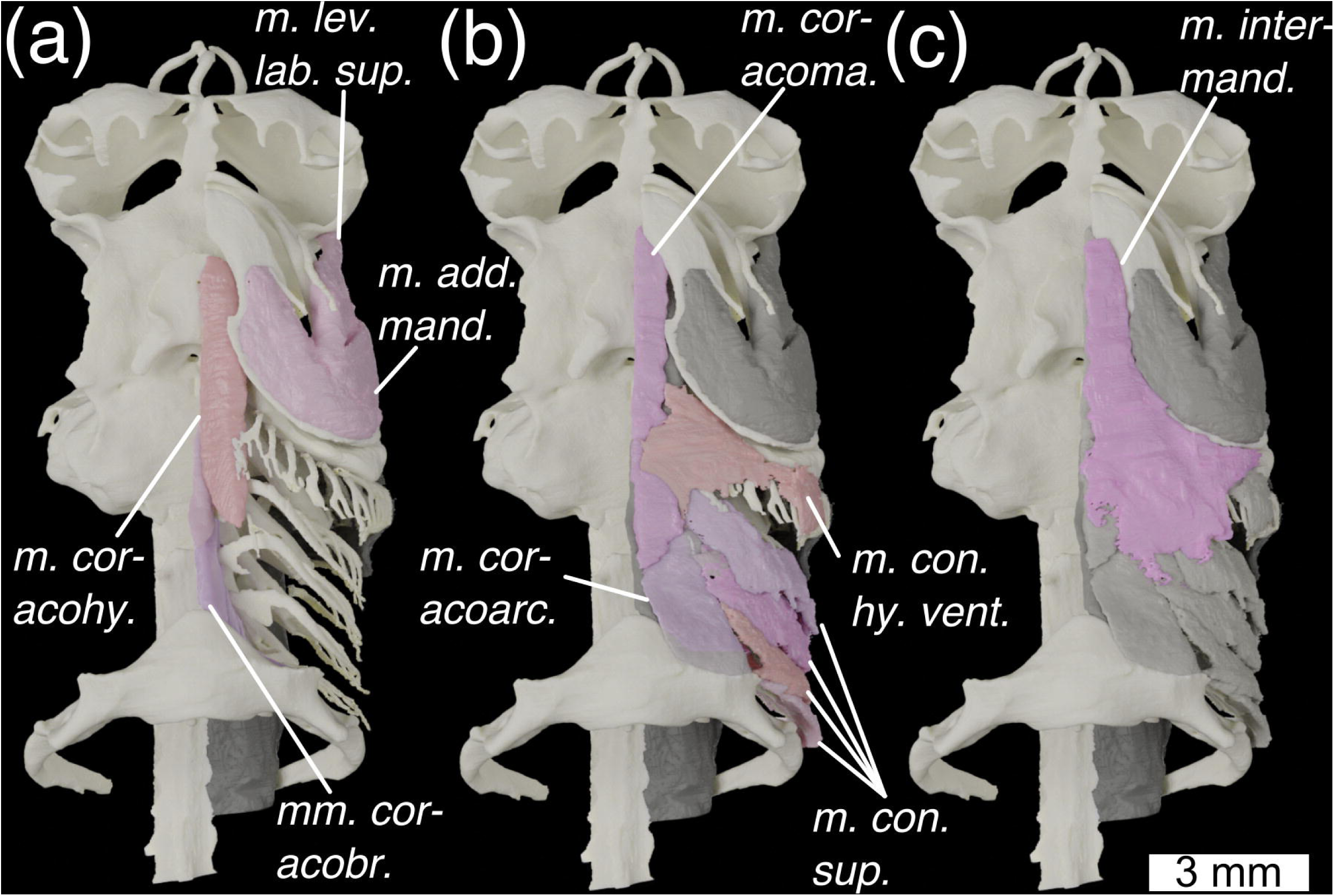
Ventral view of the head of *Scyliorhinus canicula* with (a) deepest muscles, (b) deeper muscles, and (c) shallow muscles overlain. Colours and abbreviations as in Fig. 10.

*Description:* This muscle is divided into two parts, dorsal and ventral, and is separated from the *M. levator labii superioris* by the trigeminal (V) nerve. Dorsal and ventral parts are separated by an internal fascia, latero-ventral to the mandibular joint, on which they both insert. The origin of the dorsal part occupies a shallow fossa in the posterior third of the palatoquadrate’s lateral face. The origin of the ventral part occupies a shallow fossa in the posterior half of the lateral face of Meckel’s cartilage.

*Innervation:* Trigeminal (V) nerve (Edgeworth, 1935).

***M. intermandibularis*** (Figs. 10c, 11c, 12c; *m. intermand.*)

**Figure 12.**
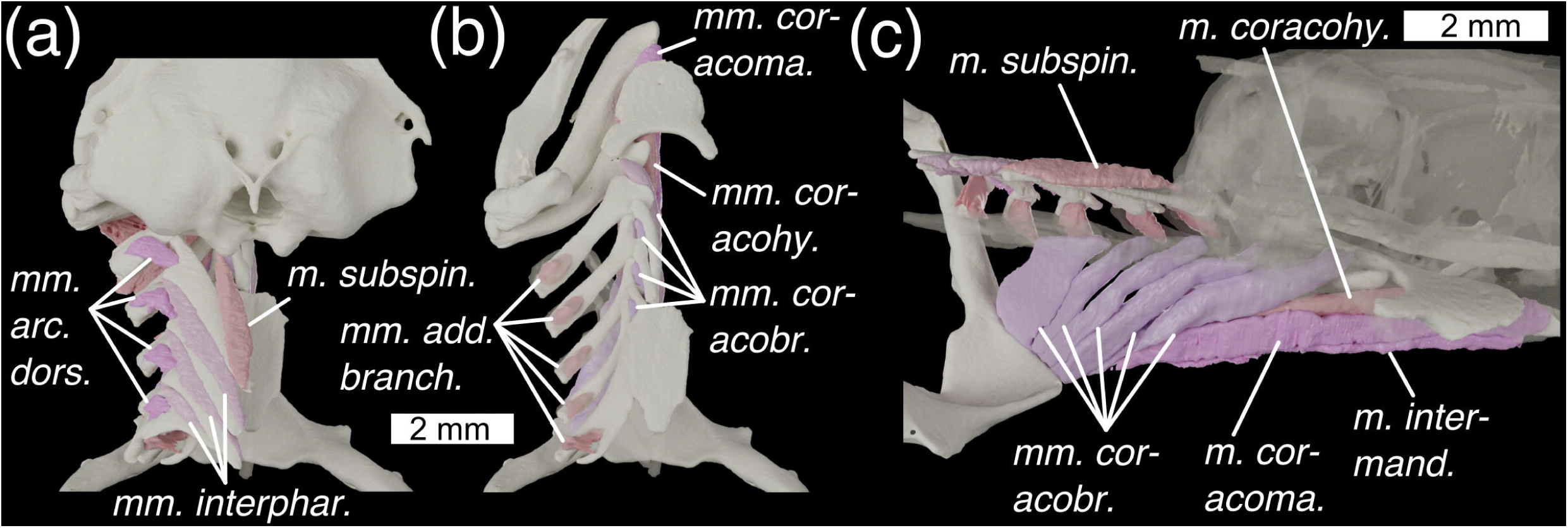
Branchial skeleton of *Scyliorhinus canicula* (a) in dorsal view, with neurocranium (b) in dorsal view with neurocranium removed and dorsal branchial skeleton semi-transparent, and (c) in medial view with nerocranium and central gill-skeleton semi-transparent. Colours and abbreviations as in Fig. 10.

*Description:* This is a broad, flat, triangular muscle with its origin along the posterior margin of the Meckelian cartilages. It then extends posteriorly as a sheet, inserting medio-ventrally on the aponeurosis of the *m. constrictor hyoideus*.

*Innervation:* Trigeminal (V) nerve (Edgeworth, 1935).

*Remarks:* The exact point of insertion is difficult to see in our scan data because it is so thin, but it matches accounts of the same muscle in other elasmobranchs (Soares and Carvalho, 2013; Ziermann et al., 2014).

***M. levator palatoquadrati*** (Fig. 10a; *m. lev. pal.*)

*Description*: A thin muscle with an origin on the latero-ventral side of the otic process, immediately posterior to the postorbital process and ventral to the neurocranial roof. It inserts on the medial side of the palatoquadrate, just anteriorly to the jaw joint, in a shallow depression.

*Innervation:* Trigeminal (V) nerve (Edgeworth, 1935).

***M. levator palpebrae nictitantis*** (Fig. 10b; *m. lev. palp. nict.*)

*Description:* A thin muscle, overlying the *m. levator palatoquadrati*. It has its origin on the on the dorso-lateral corner of the otic process, dorso-posteriorly to the origin of the *m. levator palatoquadrati* and extends antero-ventrally to insert on the lower eyelid.

*Innervation:* Trigeminal (V) nerve (Edgeworth, 1935).

***M. retractor palpebrae superioris*** (Fig. 10b; *m. ret. palp. sup.*)

*Description:* This is a short, fat muscle overlying the *m. levator palpebrae nictitantis*. It has its origin on the otic region, ventral to the origin of the *m. levator palpebrae nictitanis*. It travels dorso-anteriorly, overlying the *m. levator palpebrae nictitantis* and inserting on the upper eyelid.

*Innervation:* Trigeminal (V) nerve (Edgeworth, 1935).

*Remarks:* This matches the accounts of Ridewood (1899) and Edgeworth (1935) and is unlike data in Soares and Carvalho (2013) who imply that *Scyliorhinus* has both a *m. retractor palpebrae superioris* and a *m. depressor palpebrae superioris*.

***M. constrictor hyoideus dorsalis*** (Fig. 10c; *m. con. hy. dors.*)

*Description:* This is a large thin muscle with a broad origin extending from over the anteriormost *m. constrictor superficiales* and anteriorly over the lateral side of the *m. epaxialis*. The muscle overlies the eyelid muscles and has a further origin on the lateral wall of the otic capsule. The anterior part of the muscle inserts along the postero-ventral two thirds of the hyomandibula, while posteriorly it overlies the hyoid rays, inserting in the dorsal fibres of the *m. constrictor hyoideus ventralis*.

*Innervation:* Facial (VII) nerve (Edgeworth, 1935).

*Remarks:* This muscle includes the *m. levator hyomandibulae* (the anterior fibres), which are inseparably joined (Soares and Carvalho, 2013). Exact boundaries were difficult to segment out due to the muscle’s small width.

***M. constrictor hyoideus ventralis*** (Fig. 10c,11b; *m. con. hy. vent.*)

*Description:* This is a thin, flat muscle with a ventral origin along the medial aponeurosis, underlying the *m. coracomandibularis*. It overlies the hyoid rays and inserts into the fibres of the *m. constrictor hyoideus dorsalis*.

*Innervation:* Facial (VII) nerve (Edgeworth, 1935).

*Remarks:* Exact boundaries were difficult to segment out due to the muscle’s small width.

***M. constrictor spiracularis*** (not figured)

*Remarks:* The *m. constrictor spiracularis* lies posterior to the spiracle in *Scyliorhinus* (Ridewood, 1899). However, the constrast surrounding the spiracle was insufficient to resolve this in our scan data.

***M. adductores branchiales*** (Figs. 10a, 12b; *m. add. branch.*)

*Description:* These are small muscles of which there are five. They have their origin on spoon-shaped fossae on the dorsal surface of each ceratobranchial and insert in shallow fossae in the ventral surface of each corresponding epibranchial. The fifth inserts on the posterior pharyngobranchial complex.

*Innervation:* Vagus (X) and glossopharyngeal (IX) nerves (Edgeworth, 1935).

*Remarks:* Unlike the *Callorhinchus* there is one muscle per branchial arch.

***Mm. arcuales dorsales*** (Figs. 10a, 12a; *m. arc. dors.*)

*Description:* There are four of these muscles. They have their origin on the lateral surfaces of epibranchials I-IV, and insert on the ventro-lateral edge of each pharyngobranchial, between their anterior and lateral processes.

*Innervation:* Vagus (X) and glossopharyngeal (IX) nerves (Edgeworth, 1935).

*Remarks:* Allis (1917) refers to these as the *Mm. arcuales*. We have used Edgeworth’s name for clarity.

***Mm. constrictor superficiales*** (Figs. 10c, 11b, 12c; *mm. con, sup.*)

*Description:* These are very thin sheet-like muscles. They have their origin medially to the *m. cucullaris profundus*, where they insert between the dorsal extrabranchial cartilages. They travel ventrally to the gill bars, and insert ventrally into the medial aponeurosis.

*Innervation:* Vagus (X) and glossopharyngeal (IX) nerves (Edgeworth, 1935).

***M. subspinalis*** (Figs. 10a, 12a,c; *m. subspin.*)

*Description:* A thin flat muscle with an origin on the postero-ventral edge of the neurocranium, ventral to the occipital condyle, as well as on the ventral side of the spinal column. It then passes posteriorly and inserts on the medial tip of pharyngobranchial I.

*Innervation:* Spinal nerves, specifically the *plexus cervicalis*, formed from two or more anterior spinal nerves (Edgeworth, 1935).

***Mm. interpharyngobranchiales*** (Figs. 10a, 12a; *Mm. interphar.*)

*Description:* These pass between the pharyngobranchials and are three in number.

*Innervation:* Spinal nerves, specifically the *plexus cervicalis*, formed from two or more anterior spinal nerves (Edgeworth, 1935).

***M. coracomandibularis*** (Figs. 10a, 11b, 12b,c; *m. coracoma.*)

*Description:* This is a long thin muscle. It has its origin on the posterior part of Meckel’s cartilage, and runs posteriorly to insert with the *m. coracohyoideus* and *m. coracobranchiales* muscles ventrally (on the aponeurosis of the *m. constrictor hyoideus centralis*).

*Innervation:* Spinal nerves, specifically the *plexus cervicalis*, formed from two or more anterior spinal nerves (Edgeworth, 1935).

***M. coracohyoideus*** (Figs. 10a, 11a, 12b,c; *mm. coracohy.*)

*Description:* This is a long thin muscle. It has its origin on the ventral surface of the basihyal, and extends postero-ventrally to attach ventrally (on the aponeurosis of the *m. constrictor hyoideus centralis*).

*Innervation:* This is innervated by spinal nerves, specifically the *plexus cervicalis*, formed by two or more anterior spinal nerves (Edgeworth, 1935).

*Remarks:* Edgeworth calls this the *rectus cervicis* but for consistency we have kept it as *m. coracohyoideus*.

***Mm coracobranchiales*** (Figs. 10a, 11a, 12b,c; *mm. coracobr.*)

*Description:* These are five in number. The first has its origin anterior to the first ceratobranchial’s ventral tip. This pattern continues posteriorly with the II-IV. The Vth one has its origin on ceratobranchial V and the copula. Number I inserts at the medial part with the coracohyoid and mandibular. II-V insert on the coracoid process of the scapulacoracoid.

*Innervation:* Spinal nerves, specifically the *plexus cervicalis*, formed from two or more anterior spinal nerves (Edgeworth, 1935).

***M. cucullaris profundus*** (Fig. 10b; *m. cuc. prof.*)

*Description:* is a large triangular muscle divided into two parts by an internal septum. The anterior part has its origin in the anterior musculature, between the hyoid constricor and the epaxial muscles. It inserts on the posteriormost epibranchial. The posterior part inserts along the length of the scapular process.

*Innervation: Vagus* (X) nerve (Edgeworth, 1935).

***M. coracoarcualis*** (Figs. 10b, 11b; *m. coracoarc.*)

*Description:* A short broad muscle with an origin on the anterior face of the ventral symphysis of the shoulder girdle. It inserts on the ventral aponeurosis, between the *m. coracohyoideus* and the *m. coracobranchiales*.

*Innervation:* Spinal nerves (Edgeworth, 1935).

***M. epaxialis*** (Figs. 10b; *m. epax.*)

*Description:* Segmented muscles with an origin on the posterior part of the dorsal surface of the neurocranium, in broad fossae over the otic capsule. It inserts posteriorly with the dorsal myomeres.

*Innervation:* Spinal nerves (Edgeworth, 1935).

***M. rectus dorsalis*** (Fig. 9b; *m. rect. dors.*)

*Description:* This is antagonistic to the *M. rectus ventralis.* It has its origin in the posterior part of the orbit, antero-dorsally to the foramen for the V+VII nerves and below the entry for the superficial ophthalmic complex. It inserts dorsally over the eye.

*Innervation:* Oculomotor (III) nerve (Edgeworth, 1935).

***M. rectus ventralis*** (Fig. 9b; *m. rect. vent.*)

*Description:* This is antagonistic to the *M. rectus dorsalis*. It has its origin immediately posterior to the former’s origin, above the foramen for the V+VII nerves. It inserts ventrally around the eye.

*Innervation:* Oculomotor (III) nerve (Edgeworth, 1935).

***M. rectus lateralis*** (Fig. 9b; *m. rect. lat.*)

*Description:* This is antagonistic to the *M. rectus medialis.* It has its origin in the orbit ventrally to that of the *M. rectus ventralis* trigeminal nerve entrance. It inserts posteriorly around the eye.

*Innervation:* Abducens (VI) nerve (Edgeworth, 1935).

***M. rectus medialis*** (Fig. 9b; *m. rect. med.*)

*Description:* This has its origin relatively anteriorly to the other rectus muscles and is antagonistic to the *M. rectus lateralis.* It inserts anteriorly around the eye.

*Innervation:* Oculomotor (III) nerve (Edgeworth, 1935).

***M. obliquus ventralis*** (Fig. 9b; *m. obl. vent.*)

*Description:* This is antagonistic to the *M. obliquus dorsalis*. It has its origin anterior to the foramen for the IV nerve. It inserts antero-ventrally around the eye.

*Innervation:* Oculomotor (III) nerve (Edgeworth, 1935).

***M. obliquus dorsalis*** (Fig. 9b; *m. obl. dors.*)

*Description:* This is antagonistic to the *M. obliquus ventralis*. It has its origin just dorsal to that for the *M. obliquus ventralis*. It inserts antero-dorsally around the eye.

*Innervation:* Trochlear (IV) nerve (Edgeworth, 1935).

#### Ligaments

**Ethmopalatine ligament** (Fig. 8a; *lig. ethpal.*)

*Description:* This ligament links the palatoquadrate to the neurocranium. It attaches onto the posterior part of the nasal capsule and the anterior part of the suborbital shelf. It extends to attach on the anterior side of the ethmoid process of the palatoquadrate.

**Mandibulohyoid ligament (**not figured)

In elasmobranchs a ligament binds the hyoid arch to the mandible (Wilga *et al*. 2000). However, contrast is insufficient to model this out in our scan data.

**Superior and inferior spiracular ligaments (**not figured)

In *Scyliorhinus* two ligaments associated with the spiracle link the ceratohyal and hyomandibula to the neurocranium (Ridewood, 1896). However, the contrast around the spiracle is insufficient to model these in our data.

## Discussion

The very different cranial constructions of *Callorhinchus* and *Scyliorhinus* are reflected in two very different arrangements of cranial muscles. In *Callorhinchus* and other holocephalans, muscles are arranged to accommodate an anteriorly placed mandible, extensive labial cartilages, and a subcranial pharynx (Didier, 1995). In *Scyliorhinus* and other elasmobranchs they are instead arranged around a hyostylic jaw suspension and elongated pharynx, with some variation within the group, particularly in the areas of the eyelid muscles and jaws(Soares and Carvalho, 2013). The evolution of these two divergent morphologies can be traced back to the origins of the two clades in the Paleozoic, using the fossil record of early members of crown-group and stem-group Chondrichthyes, which preserve evidence of muscle morphology and attachments in the form of skeletal correlates and (very rarely) fossilised muscles. Assessing the relative evolution of these two types is somewhat hindered by the poorly understood phylogenetic relationships of early members of the chondrichthyan crown- and stem-groups. Nonetheless, below we assess the relationships of muscles in the living taxa described above to those seen in their fossil relatives.

### The evolution of chondrichthyan jaw musculature

Chondrichthyan jaw suspensions, around which jaw muscles are arranged, show a wide array of anatomies (Maisey, 1980), all of which likely derive from a common form seen in Paleozoic sharks. Variations in suspensory anatomy center around the palatoquadrate’s points of articulation with the neurocranium as well as the role of the hyoid arch in suspension. A prevalent idea amongst early vertebrate workers was that all gnathostome jaw suspensions derive from autodiastyly: an arrangement where the palatoquadrate is suspended from the neurocranium by otic and basal articulations and the hyomandibula is non-suspensory, a form from which in theory all gnathostome jaw suspensions can be derived (De Beer and Moy-Thomas, 1935; Grogan et al., 1999; Grogan and Lund, 2000). Autodiastyly was originally hypothetical and does not exist in any living vertebrate, although comparisons have been drawn with holocephalan embryos (Grogan et al., 1999) and a comparable arrangement exists in the Paleozoic stem-holocephalan *Debeerius* (Grogan and Lund, 2000). However, fossil morphologies in current phylogenetic topologies do not support the idea that this is the chondrichthyan ancestral state. Maisey (1980, 2001, 2008) argued that an autodiastylic chondrichthyan common ancestor was unlikely based on the prevalence of suspensory hyoid arches across the gnathostome crown-group, as well as the suspensory hyoid arch and postorbital articulation of the palatoquadrate in *Pucapampella*, likely a stem-chondrichthyan. Since then this position has been strengthened as the chondrichthyan stem-group has been populated by several other taxa, including *Gladbachus*, *Doliodus*, and *Acanthodes*, all of which possess a postorbital articulation and a suspensory hyoid arch (Maisey et al., 2009; Brazeau and de Winter, 2015; Coates et al., 2018). The identification of symmoriiformes, which also have a hyomandibula articulating on the neurocranium and a postorbital articulation of the palatoquadrate, as stem-holocephalans further supports this (Coates et al., 2017), although the possibly reduced role of the hyoid arch in jaw suspension may be a precursor to the holocephalan state (Pradel et al., 2014). All this evidence points towards the holocephalan jaw suspension being derived from an ancestral state with ethmoid and postorbital articulation in what Maisey (2008) terms archaeostyly.

As a result of this the divergent mandibular adductor musculature morphologies of holocephalans and elasmobranchs presumably derives from a more elasmobranch-like model. In *Callorhinchus* and other holocephalans the mandibular adductors are located almost entirely preorbitally, directly linking the neurocranium and the lower jaw (Fig. 3; Didier, 1995). In elasmobranchs they instead lie post/suborbitally and attach to the mandibular cartilages laterally. Anderson (2008) notes that the holocephalan condition has similarities with that of the osteichthyan *Amia*, with a neurocranial origin of the mandibular adductors and an insertion on tissue slung around the ventral side of the mandible. However, in chondrichthyans this is certainly the more derived of the two conditions: stem-chondrichthyans such as *Acanthodes* and *Gladbachus*, as well as early crown-chondrichthyans such as *Tristychius* and *Akmonistion*, lack an anteriorly restricted mandible, and possess clear fossae for a mandibular adductor spanning the lateral sides of the mandibular cartilages as in elasmobranchs, and dorsally restricted by rims precluding attachment on the neurocranium (Coates et al., 2019, 2018; Coates and Sequeira, 2001; Miles, 1973). An Upper Devonian cladoselachian preserving palatoquadrates and parts of this mandibular adductor is also consistent with this morphology (Maisey, 1989). Based on the broad presence of this arrangement across chondrichthyan phylogeny the holocephalan condition can only be interpreted as derived.

Labial muscles are present in both holocephalans and elasmobranchs, but the homologies between the two are uncertain. In holocephalans labial muscles take the form of the suite of preorbital muscles inserting on the labial cartilages: the *mm. anguli oris* and the *m. labialis*. In elasmobranchs they comprise the *m. levator labii superioris* (or *preorbitalis*), which links the palatoquadrate to the neurocranium anteriorly, and which varies in its attachment to the neurocranium in different groups (Wilga, 2005). Anderson (2008) argued that these muscles are homologous on the basis that they are both preorbital facial muscles with a trigeminal (V) innervation that insert, at least in part, on the mandible (although we found no evidence for such an insertion in *Callorhinchus*). Unfortunately, corroborating evidence in stem-group and early crown-group chondrichthyans is limited. Although a labial muscle has been reconstructed in *Acanthodes* and *Cladodus* (Lauder, 1980), this is not on the basis of any positive fossil evidence — the muscle leaves no clear skeletal attachment areas, and is not observable in cladoselachian specimens preserving muscles (Maisey, 2007, 1989). If they are homologous, the shift to a holocephalan-like insertion on the labial cartilages is presumably linked to the evolution of extensive labial cartilages and holostyly, so would likely have taken place at some point in the stem-group after the divergence of symmoriiforms, However, it remains difficult to see a clear skeletal correlate.

However, the morphology of fossil holocephalan neurocrania does provide some clues and constraints on when their preorbital mandibular muscle morphology arose. Mesozoic stem-group holocephalans – *Acanthorhina*, *Chimaeropsis, Metopacanthus, Squaloraja*, and *Isychodus –* are reconstructed with skulls with a large rostrum, shaped in such a way as to allow attachment of preorbital mandibular musculature, although their flattened skulls make this difficult to know definitively (Patterson, 1965). Moreover, the Paleozoic holocephalan *Chondrenchelys* is interpreted as having had adductor muscles that attached preorbitally on the basis of the morphology of its preorbital region (Finarelli and Coates, 2014). As Finarelli and Coates (2014) highlight, an anteriorly restricted mandible and ventral gill skeleton are not present in *Chondrenchelys*, suggesting that the preorbital attachment of mandibular adductors preceded these shifts. *Debeerius* also possesses a large rostral region, although Grogan and Lund (2000) interpret the adductor muscles as having attached on the posterior part of the palatoquadrate. Providing a lower boundary on the morphology are the more shark-like stem-holocephalans such as symmoriiformes, which have neurocrania and jaws incompatible with a crown-holocephalan-like configuration (see above). Several edestids have enlarged rostra (most obviously *Ornithoprion*; Zangerl, 1966), but also separate palatoquadrates with posteriorly located adductor muscles (see *Helicoprion*; Ramsay et al., 2015). This suggests that the development of a large ethmoid region in the holocephalan stem-group preceded the fusion of the palatoquadrates with the neurocranium and the shift of the adductor muscles onto the neurocranium, which took place somewhere between the divergence of edestids and *Chondrenchelys* from the crown-lineage.

A curious exception that does not fit this pattern is iniopterygians, another member of the holocephalan stem-group. Like in living holocephalans this group possesses anteriorly placed Meckel’s cartilages with a medial symphysis, palatoquadrates fused to the neurocranium (in some members), and a subcranial branchial skeleton (Pradel, 2010; Pradel et al., 2009). As three-dimensional specimens of “*Iniopera*” show, the large orbits would have obstructed a postorbital attachment of the adductor muscles (Pradel, 2010; Pradel et al., 2009). However, a large rostral surface for a preorbital attachment of the adductor muscles is decidedly absent. Instead a very shallow fossa above the quadrate articulation in the suborbital shelf provides a possible attachment point (Pradel, 2010; fig. 6,11), as does the small preorbital surface (Pradel, 2010; fig. 6). If two mandibular adductors were present, as in *Callorhinchus*, it is possible that the fossa in the orbit provided attachment for the posterior mandibular adductor while the anterior mandibular adductor attached preorbitally. Notably “*Iniopera*” also possesses ventrally delimited fossae on the postero-lateral sides of the Meckel’s cartilages, possibly to house the ventral adductor (Pradel, 2010; figs. 31, fam), suggesting the muscles were not slung ventrally around the jaw as in living holocephalans. Whether this set of traits is homologous with crown-holocephalans or a homoplasy linked to iniopterygians derived form is unclear. However, iniopterygians do demonstrate that once holostyly and anteriorly-placed adductor muscles had evolved, a large-rostrumed holocephalan-like form was not an inevitability for holostylic holocephalans.

As well as the facial muscles, the large coracomandibularis is an unusual feature of the holocephalan mandibular musculature. In both holocephalans and elasmobranchs, the lower jaw is depressed by the coracomandibularis muscle, unlike in osteichthyans which use the coracohyoideus to transmit movement through the mandibulohyoid ligament (Anderson, 2008). A chondrichthyan-like arrangement was also present in mandibulate stem-gnathostomes, and so is likely the ancestral state for crown-group gnathostomes (Johanson, 2003). However, within chondrichthyans the coracomandibularis of holocephalans (e.g. *Callorhinchus*, Fig. 4) is far larger than that of *Scyliorhinus* and other chondrichthyans and attaches over a greater area both on the shoulder girdle and on the mandible. Again, evidence for the presence of this muscle is limited in the holocephalan stem-group, but it seems likely to be part of the suite of adaptations linked to durophagy and a ventrally located pharynx. In the stem-holocephalan *Iniopera* the shoulder girdle and mandible have an extremely close association (Pradel pers. obs.) possibly placing the evolution of a large, short coracomandibularis as far back in the holocephalan stem-group as crown-holocephalans’ divergence from iniopterygians.

### Pharyngeal muscles

The non-suspensory nature of the holocephalans hyoid arch as well as its supposed “morphologically-complete” nature has been used to argue that they represent the primitive gnathostome condition (Grogan and Lund, 2000), but both are likely derived. Maisey (1984) outlined why the holocephalan condition is unlikely to be plesiomorphic for chondrichthyans on the basis of a convincing set of anatomical arguments. This included the fact that the holocephalan hyoid arch is bypassed by the branchial muscles linking the dorsal branchial series, the condition we see in *Callorhinchus*. In stem-chondrichthyans such as *Gladbachus* and *Acanthodes* the hyomandibula articulates directly with the neurocranium and lacks a pharyngohyal (Brazeau and de Winter, 2015; Coates et al., 2018), and the same state is seen in shark-like putative stem-holocephalans (e.g. *Ozarcus*; Pradel et al., 2014) strongly suggesting that this condition is plesiomorphic for the chondrichthyan crown-group. This implies that hypotheses based on the idea that the holocephalan branchial skeleton and jaw articulation are primitive are incorrect, and it seems likely that a more elasmobranch-like arrangement is primitive for chondrichthyans. If autapomorphic the holocephalan pharyngohyal is perhaps instead linked to the holocephalan hyoid arch’s role in supporting the roof of the pharynx, or with the role it plays in supporting the operculum. The ligament we report linking the pharyngohyal to the basicranium may anchor the hyoid arch to the neurocranium for one (or both) of these purposes.

Although both holocephalans and elasmobranchs possess a *coracomandibularis*, linking mandible and shoulder girdle, holocephalans alone possess a *mandibulohyoideus* (*interhyoideus* of Didier (1995), *geniohyoideus* of Kesteven (1933)) linking the mandible to the ceratohyal. Although morphologically similar to the *geniohyoideus* of osteichthyans, Anderson (2008) argues that the muscles are not homologous. Rather, the plesiomorphic means of depressing the mandible in crown-gnathostomes is broadly thought to be achieved by the coracomandibularis (Anderson, 2008; Johanson, 2003, p. 20; Wilga et al., 2000). In living holocephalans like *Callorhinchus* the muscle attaches to a pronounced ventral process on the broad ceratohyal. *Iniopera* also possesses a large, flat ceratohyal with a similar process, suggesting that this morphology was present at least in iniopterygians in the holocephalan stem-group (Pradel pers. obs.). *Ozarcus*, as well as stem-chondrichthyans such as *Acanthodes*, *Ptomacanthus*, and *Gladbachus* instead have long, thin ceratohyals with no such process (Coates et al., 2018; Dearden et al., 2019; Miles, 1973; Pradel et al., 2014) bolstering the idea that such a muscle was absent in the early holocephalan and chondrichthyan stem-groups, and that its evolution is linked to a holostylic jaw suspension. However, it is difficult to rule out the possibility completely: in osteichthyans with a *geniohyoideus*, for example *Amia* (Allis, 1897; Anderson, 2008) the ceratohyal is not necessarily long and thin. If this muscle is novel in chimaeroids it may be linked to their unusual ventilatory process, which is based on fore-aft movements (Dean et al., 2012) and which may have been present some chondrichthyan stem-group members (Pradel pers. obs.).

The extended and compact pharynxes in elasmobranchs and holocephalans respectively necessitate major differences in branchial musculature. Although a sub-cranial pharynx is likely plesiomorphic for crown-group gnathostomes and is present in some stem-chondrichthyans (Dearden et al., 2019), it does seem that a posteriorly extended pharynx was present at the divergence of elasmobranchs and holocephalans, given this state in several stem-chondrichthyans and putative stem-holocephalans (Coates et al., 2018; Pradel et al., 2014). Despite the major shift to a holocephalan condition, many of the muscles of the elasmobranch pharynx are preserved in *Callorhinchus* including hyoid and branchial constrictors, the *subspinalis* muscle, *interpharyngobranchialis* muscles and the *interarcuales* muscles, suggesting that these were present in the ancestral crown-chondrichthyan. Linked to this shift is the close muscular association of the neurocranium with the scapulocoracoid via the *cucullaris superficialis, cucullaris profundus,* and *protractor dorsalis pectoralis*. Skeletal correlates for these muscles are difficult to find, but a sub-cranial branchial skeleton is present in several Paleozoic stem-holocephalans including iniopterygians and *Debeerius*, suggesting that they may have been present (Grogan and Lund, 2000; Pradel et al., 2010). In *Chondrenchelys* the branchial skeleton appears to be more posteriorly placed (Finarelli and Coates, 2014). Given this and the extended pharynxes of some stem-chondrichthyans and stem-holocephalans an elasmobranch-like extended cucullaris seems likely to be plesiomorphic for chondrichthyans with a separate *cucullaris profundus* and *cucullaris superficialis* muscles having evolved in the holocephalan stem-group perhaps crownwards relative to *Chondrenchelys*.

A holocephalan-like hyoid operculum with its greatly enlarged hyoid constrictor muscles seems likely to be derived and linked to a ventrally placed branchial skeleton. However, its presence/absence can be difficult to detect in the fossil record as the evidence for gill slits/operculae is often indirect. In one Upper Devonian cladoselachian specimen the muscles and branchial rays are preserved, demonstrating an elasmobranch-like arrangement of both (Maisey, 1989). In several other Paleozoic sharks” posteriorly extended branchial skeletons such as *Triodus* and *Tristychius* “operculae have been inferred on the basis of enlarged hyoid rays, although some of these have since been disputed (discussed in Coates *et al.,* 2018, supp. mat.). An enlarged hyoid operculum is not necessarily mutually exclusive with posteriorly extending branchial arches. In several acanthodian-grade stem-chondrichthyans, where the gill slits are observable in the patterning of the dermal shagreen, both an enlarged hyoid operculum and several posterior gill slits can be observed (Watson, 1937). So, while the holocephalan sub-neurocranial pharynx is derived an enlarged operculum may run deeper into the holocephalan, or even chondrichthyan, stem-group.

### Conclusions

Here, we completely describe for the first time in 3D the two different chondrichthyan cranial muscle arrangements, in a model holocephalans and elasmobranch. The reduced, but recognizably shark-like branchial musculature we identify in *Callorhinchus*, and the likely derived origins of most holocephalan cranial musculature based on fossil evidence, do not match the idea that holocephalans display a primitive collection of archetypal chondrichthyan conditions. Instead, they fit into an emerging picture where holocephalan anatomy comprises a set of apomorphic conditions with their origins in a shark-like form (Coates *et al*. 2018). Parts of this holocephalan anatomical suite, such as the unique hyoid arch morphology and *mandibulohyoideus*, may be functionally linked to their novel mode of respiration (Dean et al. 2012). A shark-like cranial musculature, with an extended pharynx and a postorbital and ethmoid jaw suspension, seems the likely ancestral state at the chondrichthyan crown-node. Digital dissections provide a unique way of describing and sharing anatomical structures in 3D and can help identify muscles which are difficult to view in gross dissection. However, this should be seen as a complement, rather than a substitute, to traditional methods as the resolution of scan data and the inability to manipulate and examine tissues directly create limitations on what can be seen. We hope this data will, alongside digital dissections from a broad array of organisms, help others to make better inferences on the origin and evolution of the vertebrate head.

## Acknowledgments

We thank the ID-19 beamline at the European Synchrotron Radiation Facility for assistance with scanning. Assistance in getting fresh *Scyliorhinus canicula* specimens from Didier Casane and Véronique Borday-Birraux from the Laboratoire Evolution Génomes Comportement Ecologie - CNRS, and Patrick Laurenti (Laboratoire Interdisciplinaire des Energies de Demain; Université Paris-Diderot) is gratefully acknowledged. We thank Florent Goussard (CR2P - MNHN) for his inestimable help in 3D work. Some of this work began at the AMNH with the help of John Maisey.

## Author contributions

RPD, AH, and AP conceived and designed the study. PT and AP scanned the specimens. RPD, RM, AC, and AP processed the scan data. RPD analysed the data, with input from AP, AH, and DD. RPD wrote the paper and made figures. All authors read and provided feedback on the manuscript.

## Data Availability Statement

All 3D data is openly available in a public repository that issues datasets with DOIs (on publication). Tomographic data is available on request from the authors.

## Supporting Information

The datasets supporting this work are available at the private figshare links below. Included are 3D pdfs and plys of all models.

**SI1:** 3D pdfs, https://figshare.com/s/1e16cd81293498b970b5

**SI2:** *Callorhinchus* 3D models, (plys): https://figshare.com/s/6e5b73cc5b65790f2a33

**SI3:** *Scyliorhinus* 3D models, (plys): https://figshare.com/s/815ceadc2a356d9ec79d

